# Harnessing catalytic knowledge to guide enzyme discovery and evolution with AlphaEnzyme

**DOI:** 10.64898/2026.11.04.764416

**Authors:** Yong Liu, Jianxin Liu, Zhanpeng Shi, Renjian Song, Ziyan Zhao, Beibei He, Xiaohui Liu, Hongyan Liu, Jing Chen, Kaiqi Li, Jiahao Wang, Yanjie Huang, Junxi Wang, Yizhou Dang, Chang-Ting Chen, Shuangjia Zheng

## Abstract

Retrieving an enzyme for a target reaction provides a starting point for evolution, but predicting how to improve it remains difficult. This requires connecting evolutionary information with the structural and chemical context of catalysis. Here we present AlphaEnzyme, an enzyme foundation model built on a structure-grounded representation of an enzyme paired with its reaction, from which catalytic compatibility, kinetics and mutational response are read out together. On a time-separated benchmark, AlphaEnzyme achieves a top-1 catalyst retrieval success rate of 30.1%, against 17.2% for the strongest baseline. Michaelis constant predictions achieve the highest rank correlations among the compared methods, reaching 0.64 for enzymes sharing at most 40% identity with the kinetic training set. The model also enriches high-fitness TrpB variants at a budget of 16 candidates without target-specific fitness training. In prospective transaminase engineering, several prioritized variants exceed the parent, with the best design yielding a 30.4-fold increase in product signal in a lysate-based assay while retaining (*R*)-selectivity. Together, AlphaEnzyme provides a common molecular basis for selecting starting enzymes and prioritizing sequence changes for their improvement.

## 1 Main

Enzymes achieve selective chemical transformations through interactions between their active sites and reacting molecules. Many also catalyse reactions beyond their established biological roles, providing starting points for developing new catalytic functions [Buller et al., 2023]. Identifying an enzyme for a target reaction is therefore an important step in biocatalyst discovery. Its usefulness also depends on its kinetic properties and the effects of subsequent sequence changes [Savile et al., 2010, Arnold, 2018]. A candidate may support the desired transformation yet perform poorly at the substrate concentrations relevant to an application. Improving it requires predicting which mutations will preserve catalytic function and which could enhance it. These questions connect enzyme retrieval with quantitative functional prediction and the selection of variants for engineering.

Protein structure and reaction chemistry provide a basis for this connection. Evolutionary sequence models capture constraints from natural diversity [Laine et al., 2019, Rives et al., 2021], while structure-aware approaches describe residue geometry and the compatibility of a sequence with its fold [Hsu et al., 2022]. Combining these sources can improve the selection of functional enzymes, although confidence in a predicted structure alone does not consistently predict catalytic activity [Johnson et al., 2024]. Substrate-aware and reaction-conditioned models extend this description by representing proteins together with the molecules or transformations they act on [Cui et al., 2025, Li et al., 2025, Mikhael et al., 2024]. Such approaches can distinguish catalytic tasks that are poorly specified by protein sequence alone and relate predictions to the intended chemistry. In parallel, kinetic modelling has moved towards combining protein and molecular representations to predict quantitative catalytic parameters, supported by curated measurements across enzyme diversity [Muir et al., 2025, Boorla and Maranas, 2025, Wang et al., 2025]. Mutation-effect predictors use evolutionary constraints and learned sequence or structural representations to rank variants, with deep mutational scanning providing training and benchmarking data [Notin et al., 2023, Yang et al., 2025]. These developments provide complementary views of enzyme function, from structural constraints and reaction compatibility to catalytic performance and sequence variation. Connecting them within a common representation could allow knowledge learned from broad functional annotations to inform quantitative properties and mutational responses.

Reaction-conditioned retrieval models, including EnzymeCAGE, have shown that structural, evolutionary and reaction information can be combined to identify proteins compatible with a specified transformation [Liu et al., 2026]. Extending this approach to enzyme evolution requires a representation that remains informative about quantitative catalytic properties and the effects of sequence changes. Protein structure offers a natural starting point because it describes how residues are organized around the substrate and connected to the surrounding fold. Structure-prediction foundation models [Jumper et al., 2021, Abramson et al., 2024] provide learned representations of protein geometry and molecular interactions. Fine-tuning structure-prediction networks or training task-specific models on their representations has enabled variant-effect and protein-property prediction [Cheng et al., 2023, Passaro et al., 2025]. An open question is how effectively these structural representations can be combined with catalysis context to support compatibility, kinetic and mutation-effect prediction.

Here we present AlphaEnzyme, a context-aware enzyme foundation model that extends reaction-conditioned retrieval to kinetic and mutation-effect prediction. Built on an AlphaFold-like structural backbone, AlphaEnzyme combines protein structure and reaction chemistry in a shared latent representation of catalytic function. Extensive benchmarks demonstrate that AlphaEnzyme improves catalyst retrieval and Michaelis constant prediction over the compared methods, while achieving competitive performance on mutation effects under low-homology holdouts. We further evaluate variant prioritization using candidate libraries drawn from a combinatorially complete TrpB landscape [Johnston et al., 2024]. The model selects high-fitness variants under a realistic screening budget without using measurements from that landscape for training or model selection. In prospective transaminase screening, the best-performing prioritized variant gives approximately 30-fold the parental product signal in a lysate-based assay while retaining (*R*)-selectivity. These results connect knowledge learned across enzyme diversity with the local sequence changes needed to improve a selected catalyst. Ultimately, AlphaEnzyme provides a practical framework for predicting catalytic function and guiding enzyme improvement. We make the data, trained models and inference code publicly available at https://github.com/GENTEL-lab/AlphaEnzyme to support reproducible use and adaptation to new enzyme systems.

## 2 Results

### 2.1 A shared structure-grounded representation connects enzyme function across scales

Biocatalytic engineering requires solving functional prediction tasks across distinct molecular scales, including identifying natural enzymes that catalyse target transformations, evaluating their catalytic kinetics, and prioritizing active-site substitutions for directed evolution (Fig. 1a). To unify these complementary tasks within a single molecular foundation model, AlphaEnzyme learns a joint, structure-grounded representation of enzymes and their cognate reactions (Fig. 1c). The architecture couples single and pairwise structural representations from an AlphaFold-style trunk with protein language model sequence features from ESM-C, differential reaction finger-prints (DRFP), and geometric descriptors of predicted catalytic pockets. Three task-specific read-outs predict catalytic compatibility, kinetic parameters (*k*_cat_ and *K_m_*), and variant fitness effects through distinct biochemical supervision. For mutational effect prediction, this shared representation is further augmented with a parent-relative sequence residual that captures local mutational shifts (Supplementary Fig. S1). We pretrained the shared representation on approximately 500,000 curated enzyme–reaction pairs from the RHEA database before introducing kinetic or mutational supervision. Unannotated enzyme–reaction pairs served as training negatives, and training sequences were constrained to 700 residues (Methods).

**Figure 1:**
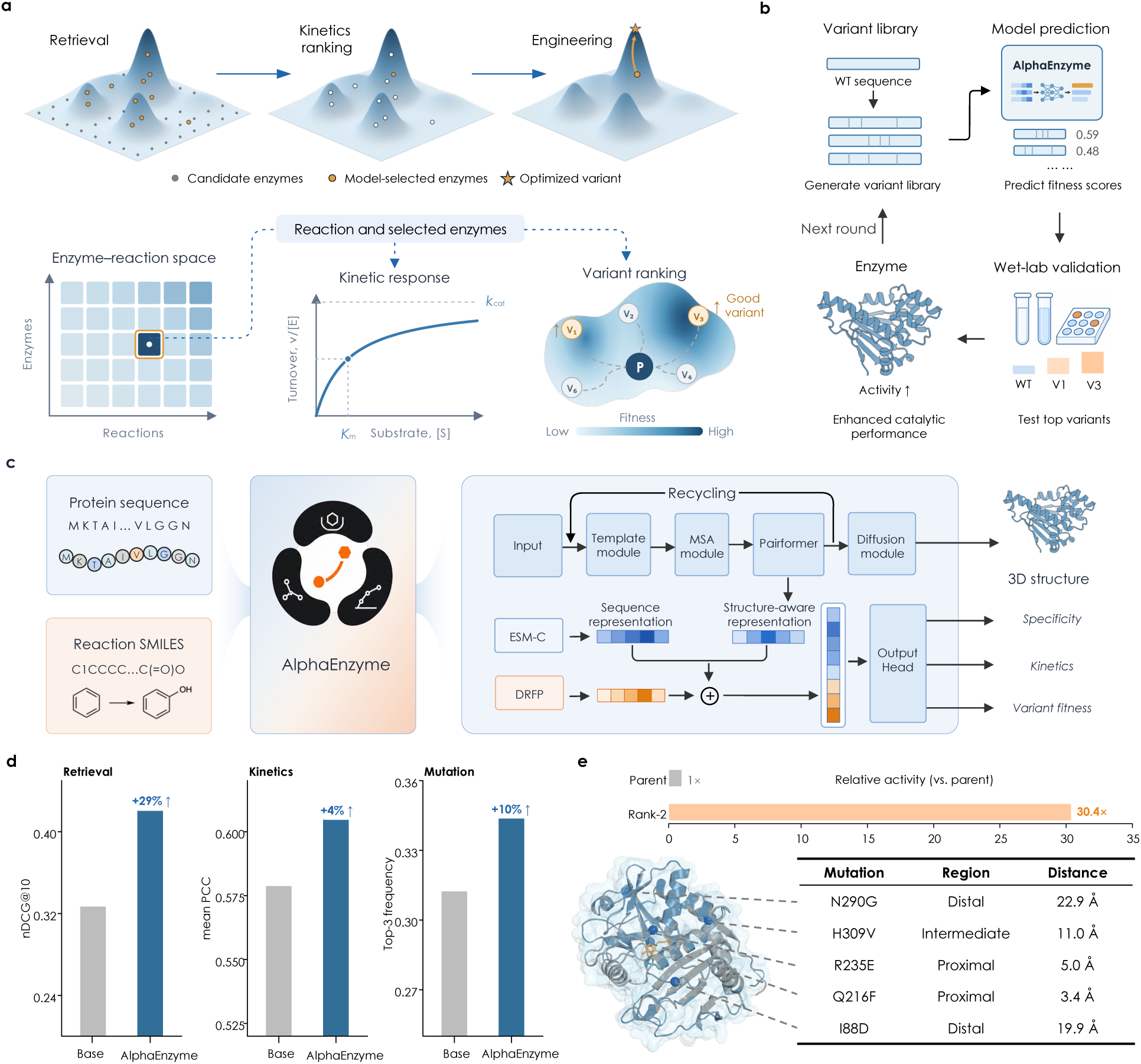
Overview of the AlphaEnzyme framework. a, Three successive enzyme engineering tasks comprising catalyst retrieval, kinetic ranking, and variant fitness scoring around parent sequence P, alongside corresponding model readouts. b, Prospective engineering cycle spanning library design, computational scoring, and experimental screening. c, Model architecture, integrating sequence representations, structural context, reaction fingerprints, and catalytic pockets into task-specific prediction heads. d, Benchmark performance across all three prediction tasks against the strongest baseline, with relative gains indicated. e, Prospective transaminase engineering round showing the rank-2 candidate, its five substitutions, and their active-site distances.

To evaluate model generalization under realistic experimental distribution shifts, each readout was tested across dedicated holdout benchmarks. These evaluations comprised a temporal partition for catalytic compatibility, sequence-identity clustering down to 40% for kinetics, and protein-level leave-one-out partitions for mutational response. Across these benchmarks, AlphaEnzyme consistently improved catalyst retrieval and Michaelis constant (*K_m_*) prediction relative to specialized baselines, while achieving competitive ranking performance across mutational landscapes (Fig. 1d). Moving beyond retrospective validation, we deployed AlphaEnzyme in a prospective engineering campaign targeting an amine transaminase (Fig. 1b). By screening only top-ranked variants from a computationally prioritized library, we isolated an engineered catalyst that exhibited approximately 30-fold the parental product signal through five amino acid substitutions (Fig. 1e).

### 2.2 AlphaEnzyme accurately identifies catalysts for unseen reactions

Catalytic compatibility defines whether an enzyme can catalyse a specific chemical transformation. AlphaEnzyme assigns a continuous compatibility score to each enzyme–reaction pair, enabling candidate enzymes to be ranked for a query reaction without reliance on Enzyme Commission (EC) numbers (Fig. 2a). We evaluated this capability on a temporally held-out test cohort derived from RHEA, spanning diverse catalytic mechanisms, enzyme classes, and sequence lengths [Bansal et al., 2022]. Using July 2023 as the pretraining cutoff, we assembled 279 reaction queries associated with 515 positive enzyme–reaction relationships deposited exclusively after this date.

**Figure 2:**
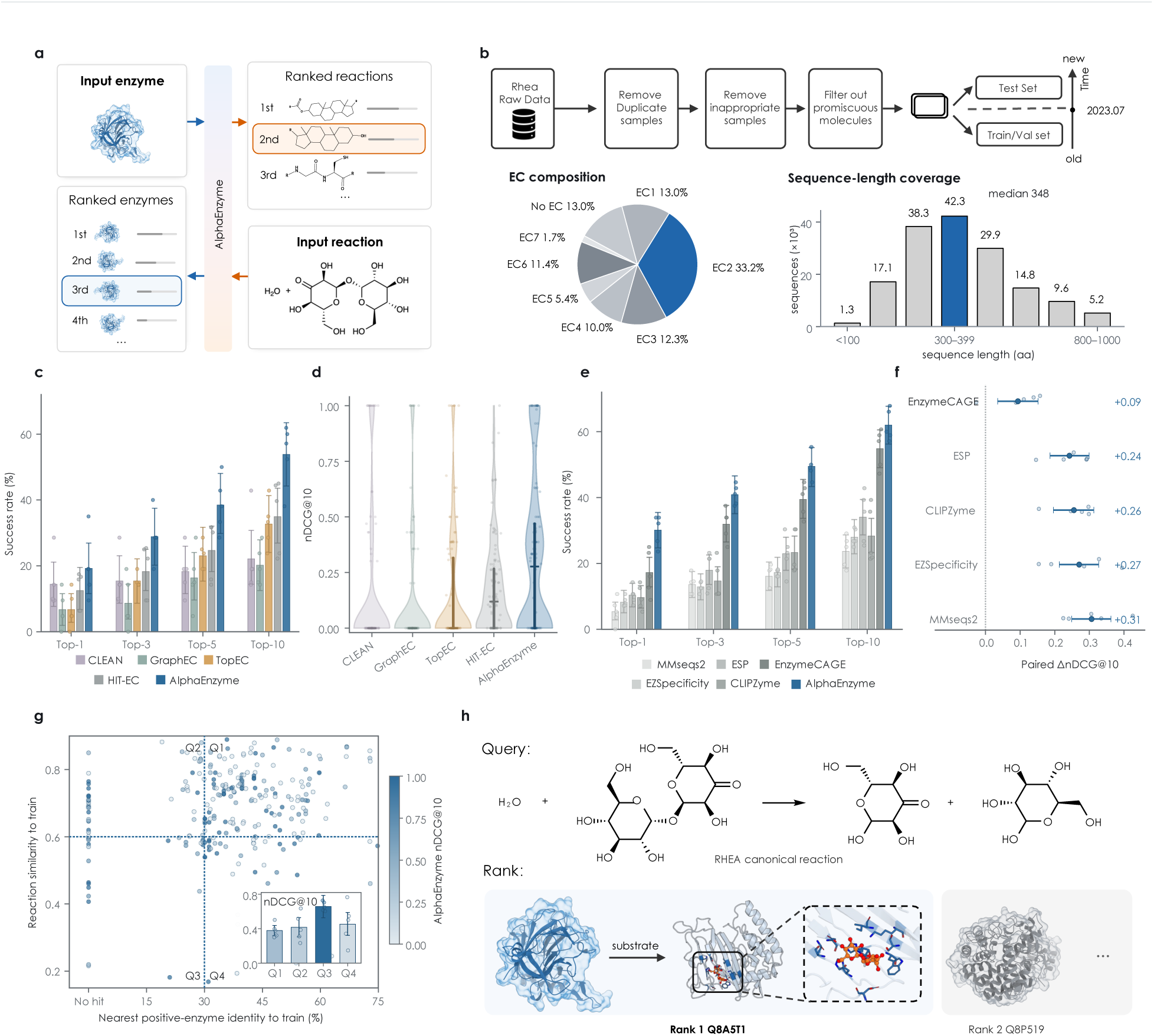
Evaluation of catalytic compatibility prediction. a, Bidirectional candidate-ranking workflow for scoring enzyme–reaction compatibility. b, Test cohort curation from RHEA, illustrating temporal separation, EC class composition, and sequence-length distributions. c,d, Top-*k* recovery and query-level nDCG@10 across 104 EC-annotated reaction queries. e,f, Retrieval performance across 279 temporally held-out queries and paired differences relative to baseline methods, where bars show query means, and points with error bars represent fold means and 95% bootstrap confidence intervals. g, Stratification of query-level nDCG@10 by reaction fingerprint similarity and sequence identity to the nearest positive enzyme, with dashed lines marking thresholds of 0.60 reaction similarity and 30% sequence identity. h, Illustrative reaction query, top-ranked catalysts, and predicted active-site enzyme–substrate complex.

None of these test annotations was used during gradient-based training (Fig. 2b, Supplementary Fig. S2a and Supplementary Table S1). Candidate pairs were drawn from the curated RHEA collection used to build the representation (Methods 4.3). Negative pairs were constructed within this collection and verified against documented positives, meaning that negative labels indicate the absence of a RHEA annotation rather than experimentally verified inactivity.

We conducted two complementary retrieval benchmarks. The first evaluated 104 reactions with complete four-digit EC annotations across 5,589 candidate pairs, including 231 known positives. Here we compared AlphaEnzyme with CLEAN, GraphEC, and TopEC, scoring these baselines by hierarchical agreement between each candidate enzyme’s predicted EC number and the query reaction’s EC number [Yu et al., 2023, Song et al., 2024, van der Weg et al., 2025]. AlphaEnzyme instead ranked candidates directly by its compatibility score without access to EC labels. The second benchmark included all 279 queries and evaluated AlphaEnzyme against the sequence-alignment method MMseqs2, the enzyme–substrate models ESP and EZSpecificity, and the reaction-conditioned retrieval models CLIPZyme and EnzymeCAGE [Steinegger and Söding, 2017, Kroll et al., 2023, Cui et al., 2025, Mikhael et al., 2024, Liu et al., 2026]. All methods within each comparison received identical candidate pools. We quantified top-*k* recovery as the proportion of queries with at least one true positive among the top *k* candidates, and normalized discounted cumulative gain at rank 10 (nDCG@10) using binary relevance, logarithmic discounting, and normalization by the ideal ranking for each query (Methods 4.4).

On the 104 EC-annotated reaction queries, AlphaEnzyme retrieved a validated catalyst among the top ten candidates in 53.8% of queries, compared with 32.7% for TopEC and 22.1% for CLEAN, achieving a mean nDCG@10 of 0.307 (Fig. 2c,d and Supplementary Table S2a). Across the broader 279-query cohort, top-ten recovery reached 62.0% compared with 54.8% for EnzymeCAGE, the strongest baseline, with a mean nDCG@10 of 0.421 versus 0.327 (Fig. 2e,f and Supplementary Table S2b). The performance advantage was most pronounced at the very top of the ranking, where top-one recovery reached 30.1% for AlphaEnzyme compared with 17.2% for EnzymeCAGE. Detailed rank distributions, enrichment factors, and top-50 recovery curves for this cohort are reported in Supplementary Fig. S3.

Because aggregate performance metrics can be driven by queries that closely resemble pretraining data, we stratified the 279 queries by reaction fingerprint similarity and sequence identity to the nearest positive catalyst (Fig. 2g, Supplementary Fig. S4 and Supplementary Table S2c,d). AlphaEnzyme outperformed EnzymeCAGE in the higher-similarity regime (162 queries, mean Δ,nDCG@10 of 0.110, 95% bootstrap confidence interval 0.036–0.186), and the margin more than doubled in the 24 jointly divergent queries where reaction similarity was at most 0.60 and positive-enzyme sequence identity was at most 30% (mean Δ,nDCG@10 of 0.265, 95% confidence interval 0.114–0.426). The two models performed comparably when only one axis diverged (Supplementary Table S2d), indicating that the advantage of the structure-grounded representation emerges once both the transformation and its candidate catalysts fall outside annotated precedent, the regime most relevant to finding enzymes for poorly characterized reactions. Absolute nDCG@10 values were higher in the jointly divergent stratum (0.657) than in the higher-similarity stratum (0.379) for four of the six evaluated methods, with ESP and MMseqs2 as exceptions. Absolute scores are not comparable across strata, because nDCG@10 is normalized within each query against its own ideal ranking, and queries in the jointly divergent stratum carry fewer annotated positives, so placing a single correct enzyme near the top already yields a high value. The paired within-stratum comparisons therefore remain the informative measure.

In an illustrative query, AlphaEnzyme ranked the known catalyst Q8A5T1 first and predicted an enzyme–substrate complex positioning the substrate within the catalytic pocket (Fig. 2h). Across both benchmarks, AlphaEnzyme recovered validated catalysts more reliably than EC-based classifiers, sequence alignment and reaction-conditioned retrieval models, with its largest margins at the top of the ranking and in the regime where the reaction and its candidate catalysts both lie outside annotated precedent, which is where retrieval accuracy determines how many candidates must be tested experimentally. Having confirmed catalyst identification for uncharacterized transformations, we next examined whether the shared representation could quantitatively predict enzymatic kinetic parameters.

### 2.3 Kinetic prediction extends across sequence-divergent enzymes

Enzymes that catalyse identical chemical transformations can exhibit substantially divergent kinetic profiles, which govern their practical suitability as industrial biocatalysts. We therefore predicted the turnover number (*k*_cat_) and Michaelis constant (*K_m_*) directly from the shared representation, grouping test enzymes by sequence clusters and systematically reducing their maximum sequence identity to any training sequence from 99% down to 40% (Fig. 3a,b).

**Figure 3:**
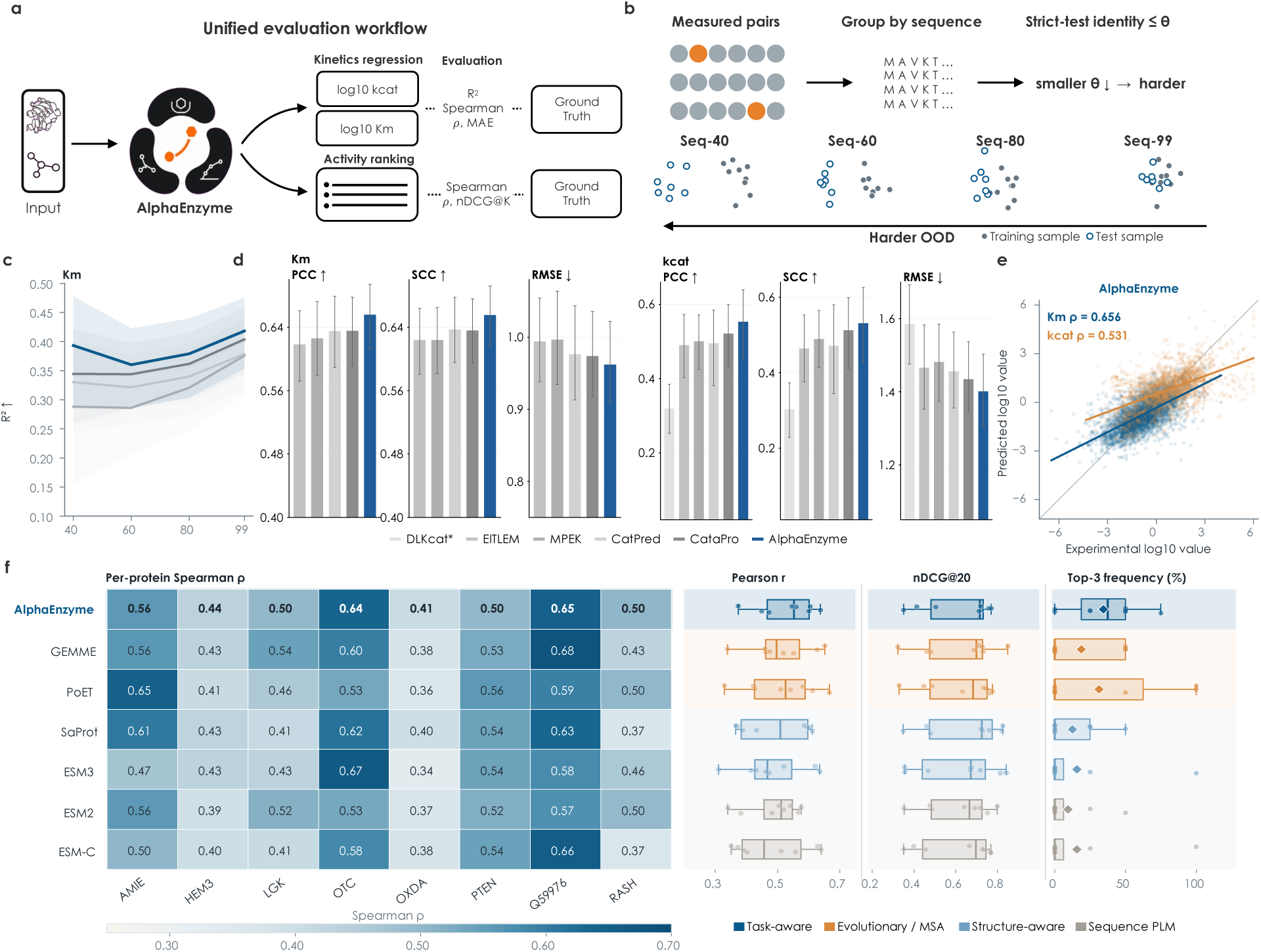
Kinetic and mutation-effect evaluation. a, Evaluation schemes for catalytic kinetics and variant fitness ranking. b, Sequence-clustered kinetic evaluation subsets, where lower identity thresholds denote greater evolutionary divergence from training enzymes. c, Fixed-checkpoint *R*^2^ values for *Km* across Seq-40, Seq-60, Seq-80, and Seq-99, with shading indicating 95% sequence-bootstrap confidence intervals. d, Pearson correlation, Spearman correlation, and root-mean-square error comparisons. e, Predicted versus experimentally measured values on the Seq-99 benchmark. f, Mutation-effect evaluation across eight leave-one-protein-out folds, where the heatmap displays per-protein Spearman correlations and box plots summarize Pearson correlations and nDCG scores. Top-3 frequency denotes the fraction of assay–metric comparisons in which a model placed among the three highest-performing methods.

The two kinetic parameters showed contrasting predictive behaviors. Prediction of *K_m_* surpassed the evaluated baselines across all sequence-divergence thresholds, with the performance advantage widening as sequence identity decreased. At the Seq-40 threshold, AlphaEnzyme achieved a Spearman correlation of 0.640 compared with 0.593 for CatPred, the highest-performing base-line at this level, yielding an *R*^2^ of 0.393 versus 0.343 and a root-mean-square error (RMSE) of 1.007 versus 1.048. In contrast, prediction of *k*_cat_ did not consistently decouple from the baseline models when sequence identity fell below 80%. At the Seq-40 threshold, AlphaEnzyme yielded a Spearman correlation of 0.391, outperforming CatPred (0.312) and CataPro (0.378) while remaining below EITLEM (0.422), with RMSE values across the five evaluated methods spanning 0.096 log units (Supplementary Table S3). AlphaEnzyme led across both kinetic readouts simultaneously on the least restrictive Seq-99 benchmark, achieving a Spearman correlation of 0.531 for *k*_cat_ and 0.656 for *K_m_* (Fig. 3d,e and Supplementary Figs. S6–S7).

This performance divergence likely reflects fundamental differences in the physical determinants of the two parameters. Active-site pocket geometry and steric complementarity provide rich structural cues for substrate recognition and apparent binding affinity, which are captured by *K_m_*. By contrast, the turnover number *k*_cat_ is governed by transition-state stabilization, conformational dynamics, and reaction-condition nuances that are less accessible from static structural representations. Consequently, while AlphaEnzyme reliably generalizes *K_m_* prediction across distant sequence homologs, its capacity to rank *k*_cat_ on unfamiliar enzymes remains comparable to specialized turnover models. The shared representation therefore transfers quantitative catalytic behaviour to enzymes far outside the training distribution, ranking *K_m_* most accurately among the compared methods down to 40% sequence identity while remaining competitive on turnover. Having established that the representation supports quantitative property prediction, we next asked whether it also captures the functional consequences of individual sequence changes.

### 2.4 Mutational knowledge transfers across enzyme families

We evaluated variant fitness ranking within individual enzyme fitness landscapes using eight enzyme deep mutational scanning assays from ProteinGym under an eight-fold leave-one-protein-out cross-validation scheme [Notin et al., 2023]. In each fold, one protein was held out for testing and one for validation, while the remaining six proteins served as training data, with performance metrics macro-averaged across the eight test enzymes. Because these eight proteins were examined during initial model development, we treat this evaluation as an exploratory benchmark rather than a blind test.

Across the eight test enzymes, AlphaEnzyme reached a mean Spearman correlation of 0.525, nominally the highest among the sixteen evaluated methods, though the margin over GEMME (0.519) and TranceptEVE (0.514) remained within fold-to-fold variation (Fig. 3f and Supplementary Table S5). The primary distinction emerged at the top of the fitness ranking, where AlphaEnzyme achieved a mean nDCG@20 of 0.622 and ranked among the top three models in 28.1% of the 32 assay–metric comparisons (Supplementary Fig. S9 and Supplementary Table S6).

Systematic ablation experiments across all three readouts delineated the specific contributions of individual model components (Supplementary Figs. S5, S8 and S9, and Supplementary Table S6). For mutational fitness, the parent-relative sequence residual provided the primary basis for local ranking, and its removal degraded every metric. For catalytic compatibility and kinetics, structural features, pocket geometry and the reaction or substrate descriptors each contributed, and removing any one of them lowered performance. Because practical protein engineering campaigns do not optimize global correlation coefficients but instead select variants from the very top of candidate rankings, we next focused on model performance under restricted experimental screening budgets.

### 2.5 AlphaEnzyme-guided selection enriches high-fitness TrpB variants

Whereas comprehensive mutational scans measure thousands of variants across a sequence landscape, practical engineering campaigns must discover rare high-fitness substitutions under strict experimental screening budgets. We therefore evaluated model-guided selection under finite candidate budgets using five disjoint libraries derived from an experimentally mapped combinatorial fitness landscape of tryptophan synthase *β*-subunit (TrpB) [Johnston et al., 2024]. Each library contained 384 combinatorial variants defined at active-site positions 183, 184, 227, and 228 around the parental VFVS sequence (Fig. 4a). No TrpB fitness measurements or related sequence data were included during training or model selection, providing an entirely unseen landscape for testing.

**Figure 4:**
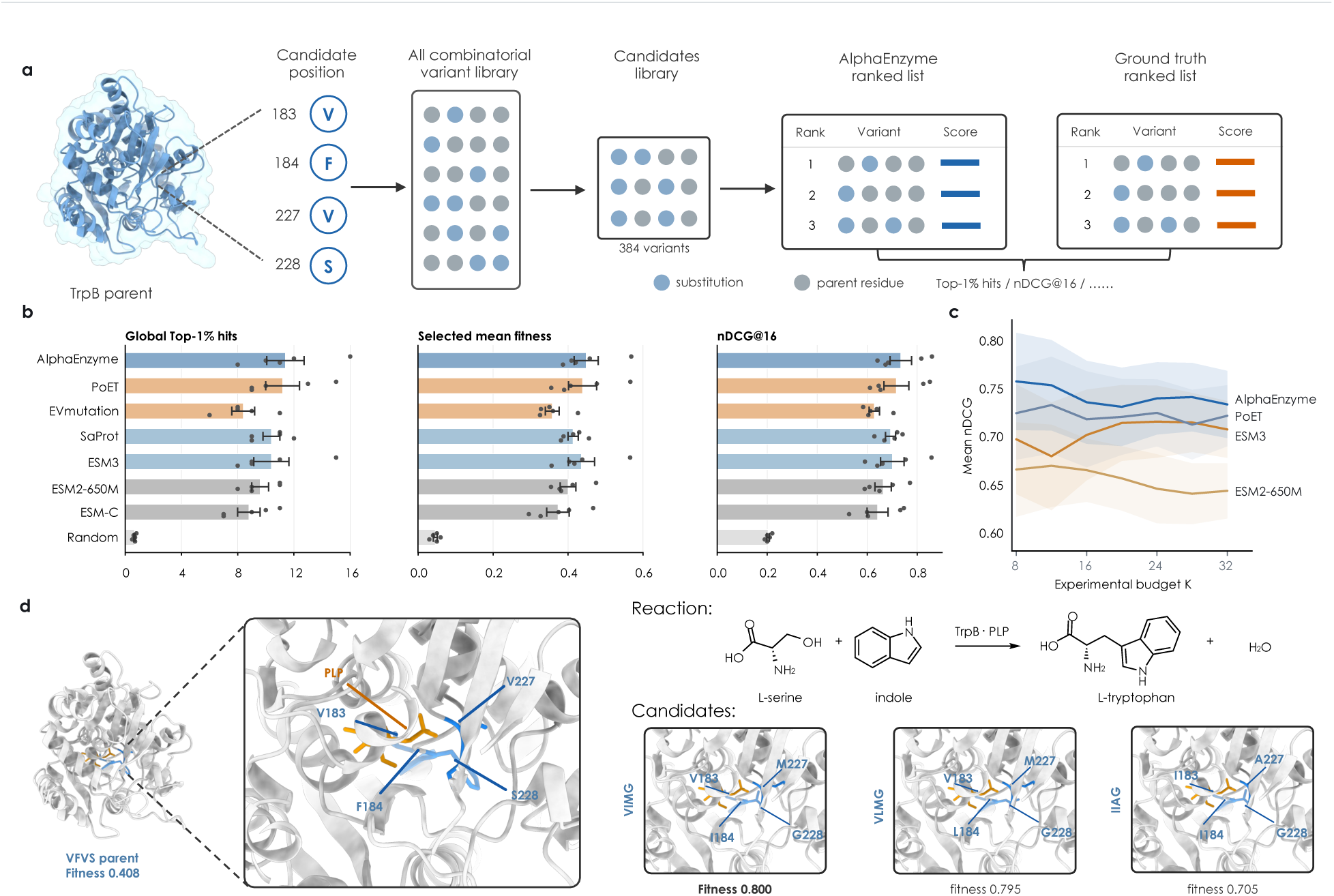
Variant prioritization in the TrpB fitness landscape. a, Model-guided prioritization of 384 combinatorial variants at positions 183, 184, 227, and 228 of the parental VFVS enzyme. b, Global top-1% hits, selected mean fitness, and nDCG@16 at a budget of 16 variants per library, where open symbols represent individual libraries, and bars with error bars denote means and standard deviations across five libraries. c, Mean nDCG across candidate selection budgets ranging from 8 to 32 variants, with shading representing standard errors of the mean across five libraries. d, Structural environment of engineered positions surrounding pyridoxal 5′-phosphate, illustrating the TrpB catalytic transformation and three prioritized variants modeled on the parent backbone.

At a screening budget of 16 variants per library, AlphaEnzyme achieved the highest cross-library average for global top-1% hits, mean selected variant fitness, and nDCG (Fig. 4b). This enrichment advantage was maintained across candidate budgets ranging from 8 to 32 variants, indicating that the performance benefit was not sensitive to a single arbitrary cutoff (Fig. 4c). Furthermore, prioritized high-fitness variants originated from multiple diverse libraries rather than being concentrated within a single panel. For instance, the variants VIMG and VLMG, each ranked fifth within their respective libraries, exhibited experimental fitness values of 0.800 and 0.795 compared with 0.408 for the parental VFVS enzyme, with VIMG achieving 1.96-fold the parental fitness. Similarly, the variant IIAG, ranked fifteenth, reached an experimental fitness of 0.705 (Fig. 4d). All prioritized substitutions mapped to the active-site pocket directly coordinating the pyridoxal 5′-phosphate (PLP) cofactor. Model-guided selection therefore enriched rare high-fitness variants on a landscape absent from training, at budgets comparable to a single round of laboratory screening and without any target-specific fitness data. Because this landscape had nonetheless been measured exhaustively in advance, we next applied the same prioritization strategy prospectively, to an enzyme and a reaction for which no fitness measurements existed.

### 2.6 AlphaEnzyme prioritizes a transaminase variant with 30.4-fold higher product signal

Amine transaminases are prominent biocatalysts for the enantioselective synthesis of optically pure chiral amines, which serve as essential building blocks for pharmaceutical synthesis [Kelly et al., 2018, Savile et al., 2010]. A prominent target is (*R*)-3-aminobutan-1-ol (RABO), one of three key intermediates in synthetic routes to the HIV integrase inhibitor dolutegravir, which an (*R*)-selective transaminase can synthesize in a single asymmetric transformation [Tang et al., 2019]. Engineering transaminases toward non-natural substrates often involves a difficult compromise between enhancing catalytic activity and maintaining stereochemical fidelity, creating an expansive combinatorial search space that benefits from model-guided candidate selection under constrained experimental budgets [Höhne et al., 2010, Yang et al., 2019]. To preserve strict stereocontrol, we began with a parental transaminase that inherently yields the required (*R*) configuration and tasked AlphaEnzyme specifically with prioritizing variants that increase catalytic activity.

To validate the platform prospectively, we deployed AlphaEnzyme in an experimental transaminase engineering campaign where variants were computationally ranked and selected prior to gene synthesis and laboratory assay. The target biotransformation converts 4-hydroxy-2-butanone to (*R*)-3-aminobutan-1-ol using isopropylamine as the amine donor (Fig. 5a,b). Ten active-site residues were selected on the basis of pocket geometry, ESM-C single-mutation compatibility, and structural-zone representation, with 16 possible amino acid states per position. From this combinatorial space, 10,000 multi-point mutants bearing two to ten substitutions were subsampled to a 1,000-member representative panel and scored relative to the parent enzyme. We experimentally screened the top 50 candidates from the AlphaEnzyme ranking, alongside 20 variants prioritized by ESM2 and 20 randomly selected variants, using a unified 96-well expression, whole-cell bio-transformation, Marfey’s derivatization, and ultra-performance liquid chromatography (UPLC) workflow (Fig. 5c).

**Figure 5:**
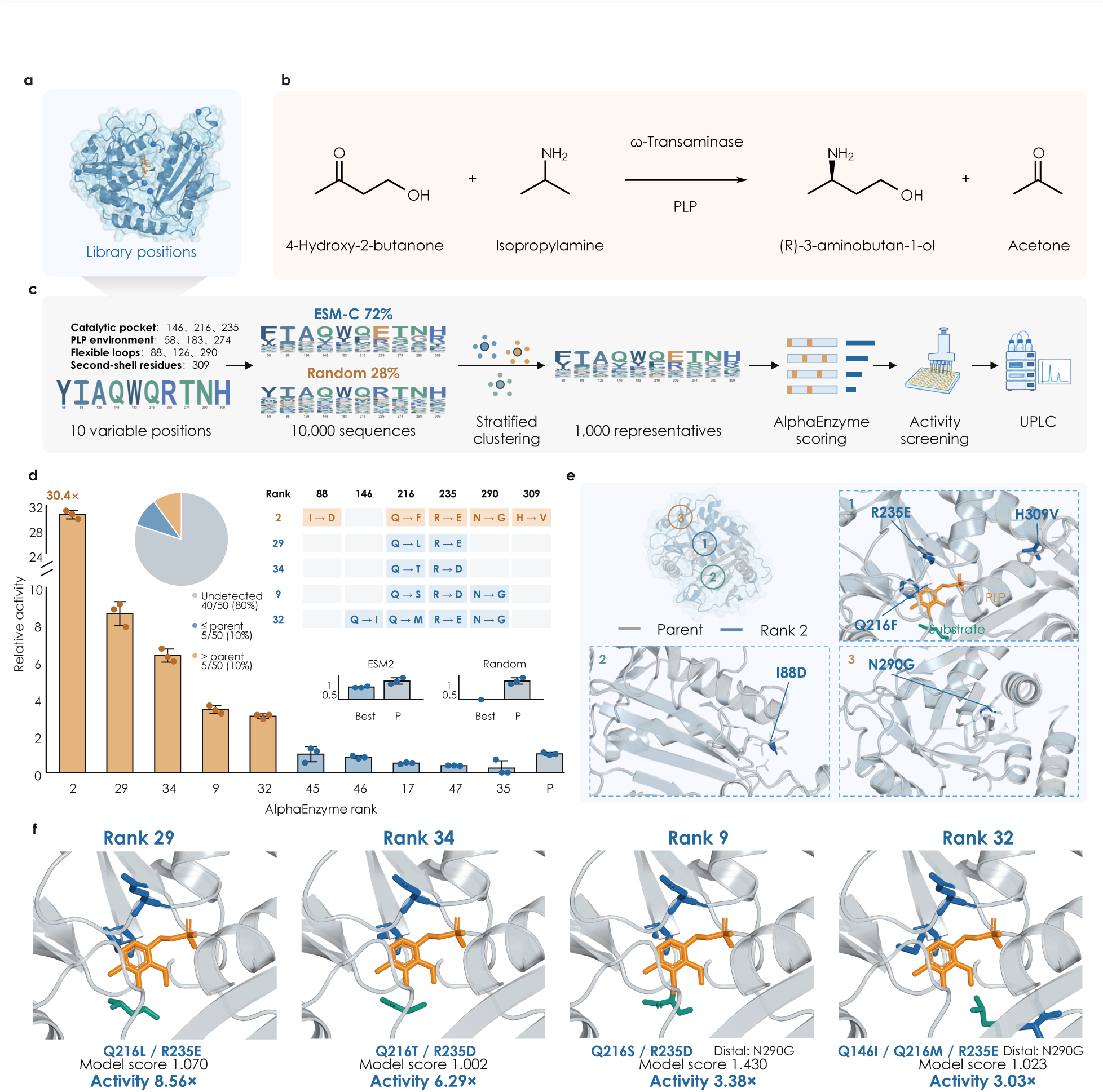
Prospective engineering of an amine transaminase. a, Predicted parental structure highlighting library-design positions in blue. b, PLP-dependent asymmetric synthesis of (*R*)-3-aminobutan-1-ol. c, Candidate library construction and screening workflow, showing proposal fractions for each stage. d, Parent-normalized RABO product signal for displayed candidates from the AlphaEnzyme top-50 cohort, ordered by measured signal and labeled by AlphaEnzyme rank, alongside same-plate parental controls (P). The embedded table lists amino acid substitutions for candidates exceeding the parental mean, and the pie chart summarizes activity outcomes across the 50 AlphaEnzyme-screened candidates. Insets depict top candidates from the ESM2-ranked and random-selection reference arms (20 candidates each) with corresponding parent controls. Bars and error bars represent means and standard deviations, points indicate three biological replicates, and parent-normalized signal denotes the chromatographic RABO peak area divided by the same-plate parent mean. e, Predicted rank-2 complex highlighting the PLP cofactor, bound substrate, and five substituted residues. f, Predicted active-site conformations for four additional candidates exceeding parental activity. All individual measurements and parental controls are available as Source Data.

The most active biocatalyst recovered from the AlphaEnzyme cohort corresponded to the candidate ranked second by the model (Fig. 5d). Its parent-normalized product signal, defined as the chromatographic RABO peak area divided by the same-plate parent mean, was 30.4 across three independent biological replicates, corresponding to a 30.4-fold increase in product formation relative to the parental enzyme under identical reaction conditions. This variant possessed five amino acid substitutions (I88D, Q216F, R235E, N290G, and H309V) relative to the parent. Four additional AlphaEnzyme candidates surpassed the parental mean, exhibiting parent-normalized signals ranging from 3.03 to 8.56 (Fig. 5d). In contrast, the top variants from the ESM2 and random selection panels failed to match parental activity, with all reference candidates remaining below the same-plate parental benchmark (Fig. 5d).

Gains in catalytic activity are practically viable only when stereochemical fidelity is preserved. Following Marfey’s derivatization, reaction products from both the parental enzyme and the rank-2 variant exclusively yielded the (*S*, *R*) diastereomeric adduct without detectable formation of the (*S*, *S*) diastereomer, confirming complete retention of the required (*R*) configuration (Supplementary Fig. S10). Structural modeling of the rank-2 complex positioned all five substitutions within the PLP-binding pocket and adjacent substrate-channel microenvironment (Fig. 5e).

Enzymatic activity in this campaign is quantified as parent-normalized product signals under standardized biotransformation conditions rather than isolated turnover constants. Under these operational conditions, a computational model trained without prior exposure to this specific enzyme, substrate, or mutational landscape placed a 30-fold improved variant at rank two and successfully recovered five improved biocatalysts from 50 tested sequences, whereas the base-line strategies produced zero improved variants across 40 candidates. A combinatorial sequence space of 10,000 variants was thus effectively navigated in a single experimental round of 90 total measurements, establishing that catalytic knowledge learned across broad enzymatic families can transfer to an uncharacterized enzyme system.

## 3 Discussion

AlphaEnzyme brings enzyme retrieval, kinetic prediction and mutation-effect prediction into a shared molecular foundation model. Building on reaction-conditioned retrieval [Mikhael et al., 2024, Liu et al., 2026], it extends the information used to identify candidate catalysts towards assessing their properties and prioritizing sequence changes.

Structure, reaction chemistry and kinetics describe related aspects of catalysis. Structure places residues in the spatial context of the catalytic pocket and surrounding fold, while reaction information specifies the transformation against which that structure must be interpreted. Kinetic measurements add quantitative information that reaction annotations alone cannot provide. Enzymes associated with the same transformation can differ substantially in their measured catalytic properties. Learning from these complementary sources could help distinguish structural features that support general protein function from those relevant to a particular catalytic task. For variant prediction, this distinction matters because a sequence change may preserve the fold yet alter substrate recognition or catalytic performance. A shared representation offers a way to connect these observations while retaining separate readouts for measurements with different meanings and experimental conditions.

We observed that the context-aware representation improved catalyst retrieval and Michaelis constant prediction and achieved competitive performance on mutation effects, consistent with complementary contributions from different sources of catalytic knowledge. In prospective transaminase screening, AlphaEnzyme identified improved variants within a limited experimental budget, whereas the ESM2-ranked and randomly selected reference groups yielded none. The best variant gave approximately 30-fold the parental product signal while retaining (*R*)-selectivity. These findings support the practical value of the model for prioritizing variants for experimental testing.

Several limitations remain. First, AlphaEnzyme scores candidate sequences rather than generating them, so its main role is in screening. This focus reflects a practical bottleneck in enzyme engineering. Many methods can propose mutations, but only a small fraction of the resulting candidates can be tested experimentally. Reliable ranking therefore remains essential, even as sequence generation improves. Second, gains in kinetic prediction were modest. Quantitative kinetic properties depend on assay conditions and other biochemical factors that are incompletely captured in current datasets, and limited measurements may further restrict predictive accuracy. Future work could integrate de novo design with catalytic scoring and incorporate richer bio-chemical and experimental context. Language model-guided workflows could help organize this information and coordinate candidate generation, prediction and experimental feedback.

Together, AlphaEnzyme links reaction compatibility, quantitative properties and variant effects through a common molecular representation. This connection provides a practical route for applying knowledge learned across enzyme diversity to the improvement of individual catalysts.

## 4 Methods

### 4.1 Task formulation

We formulate three functional readouts from a shared enzyme–reaction representation. AlphaEnzyme accepts an amino-acid sequence and a reaction represented as canonical or atom-mapped SMILES, according to the available input. Its shared encoder produces h_AE_ ∈ R^1152^. A specificity head assigns a compatibility score to each enzyme–reaction pair, permitting either member of the pair to be held fixed while the other is ranked. Separate kinetic heads estimate log_10_(*k*_cat_/s^−1^) and log_10_(*K_m_*/mM). The mutation head assigns each variant a score relative to its parent. Higher scores denote higher predicted fitness within that variant group. The reported specificity and mutation outputs are within-query ranking scores.

### 4.2 Shared structure-aware representation

The structural trunk was adapted from the AlphaFold 3-style Protenix implementation [Abramson et al., 2024, ByteDance AML AI4Science Team et al., 2025]. Sequence, template and multiple-sequence-alignment inputs are processed by the input embedder and Pairformer. For new sequences, the standard inference workflow supports a no-MSA mode. Packaged true-MSA features were retained to reproduce the frozen benchmark inputs. The diffusion module can generate a candidate structure, from which local pocket context is identified with P2Rank [Krivák and Hoksza, 2018]. For exact reproduction of benchmark predictions, we provide the frozen features used in the reported analyses. Full inference on new sequences additionally executes structure generation and pocket detection, which may exhibit minor environment-dependent numerical variation.

The shared representation comprises three projected 384-dimensional components. These encode structural single-representation features, DRFP reaction information and protein/pocket context incorporating ESM-C sequence features [ESM Team, 2024, Probst et al., 2022]. Their concatenation is

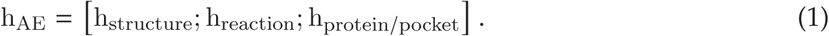

All task heads receive this 1,152-dimensional interface. The mutation workflow further uses a 1,152-dimensional sequence residual relative to the parent, as described below. Supplementary Methods M1–M3 detail the task-specific inputs and controlled readouts.

### 4.3 Specificity data and temporal separation

Experimentally annotated enzyme–reaction relationships were assembled from RHEA [Bansal et al., 2022]. Entries without a valid protein sequence or standardized reaction were removed, duplicate records were collapsed and predefined validity filters were applied. The curated collection contained 503,770 labelled pairs, comprising 478,356 training pairs and 25,414 validation pairs. These records represented 158,519 unique protein sequences and 9,624 distinct positive reactions. The active training configuration excluded proteins longer than 700 residues. Candidate negative pairs were generated within the curated collection and checked against known positive annotations.

The temporal boundary was July 2023. The temporal evaluation cohort was constructed from later RHEA annotations and remained outside gradient-based training. After canonical reaction-level deduplication, the common reaction-to-enzyme comparison contained 279 reaction queries and 515 positive relationships. Pair labels remained outside model fitting. This cohort was therefore evaluated across the annotation boundary.

### 4.4 Specificity training and evaluation

Specificity training used pairwise ranking supervision with AdamW, an initial learning rate of 5 × 10^−7^, weight decay 0.03, batch size 32 and three epochs. The schedule used a 10% warm-up followed by cosine decay, gradient clipping at 1.0 and an exponential moving average of 0.999. The interaction stack had hidden dimension 384, four transformer layers and dropout 0.1. Validation nDCG@10 was monitored during training. The resulting checkpoint was frozen before the downstream kinetic, mutation and candidate-ranking workflows.

The EC-based comparison evaluated reaction-to-enzyme retrieval on 104 reactions with complete four-level EC annotations, comprising 5,589 candidate pairs and 231 positive annotations. For each candidate enzyme, its predicted EC was compared with the target reaction EC. The ranking score was 25 times the number of matching EC fields plus the model’s native confidence. AlphaEnzyme used its native pair score. CLEAN, GraphEC, TopEC and AlphaEnzyme were evaluated on identical queries and candidates [Yu et al., 2023, Song et al., 2024, van der Weg et al., 2025]. Figure 2 also shows a contextual HIT-EC reference from three independently initialized models retrained on the EC-labelled training partition, with checkpoints selected using validation loss [Dumontet et al., 2026]. The broader 279-query temporal evaluation cohort compared AlphaEnzyme with the sequence-search method MMseqs2 [Steinegger and Söding, 2017], the enzyme–substrate models ESP and EZSpecificity [Kroll et al., 2023, Cui et al., 2025], and the reaction-conditioned retrieval models CLIPZyme and EnzymeCAGE [Mikhael et al., 2024, Liu et al., 2026]. Models within each comparison used identical query and candidate sets. Complete results are reported in Supplementary Table S2.

Top-*k* success was one for a query when at least one known positive appeared among the first *k* predictions. Discounted cumulative gain through rank 10 used binary relevance and logarithmic rank discount. This value was divided by the ideal query-specific ordering to give nDCG@10. Bars report the full-query mean. Five fixed query partitions provided the displayed fold points and visualized query-level variation. Confidence intervals were estimated with 10,000 bootstrap resamples of queries. Paired nDCG@10 differences were calculated per query before bootstrapping. For the contextual HIT-EC results, hierarchical bootstrap resampled the three training seeds and then reaction queries within each selected seed.

For the joint generalization analysis, reaction novelty was the maximum reaction-fingerprint similarity between a test reaction and the training set. Sequence novelty was the maximum sequence identity between a positive enzyme for that query and a positive training enzyme. Thresholds of 0.60 reaction similarity and 30% sequence identity defined four quadrants. Mean AlphaEnzyme nDCG@10 and 95% query-bootstrap confidence intervals were calculated within each quadrant without model retraining. Matched representation controls and resampling procedures are specified in Supplementary Methods M2 and M4. Cohort definitions are listed in Supplementary Table S1.

### 4.5 Kinetic prediction

Kinetic targets were represented as log_10_(*k*_cat_/s^−1^) and log_10_(*K_m_*/mM). Protein sequences were grouped before splitting, and strict test subsets were formed at maximum training-sequence identities of 40%, 60%, 80% and 99%. The nested subsets were evaluated with one Seq-99 training regime and checkpoint. Exact measurement counts are retained in the source data.

The kinetic path used h_AE_ as its sole predictive molecular representation, without auxiliary retrieval, EC-label, experimental-metadata, teacher-target or post hoc calibration features. Rank-4 low-rank adaptation was applied in the representation path, and a neural TabM head was trained separately for each target. CatPred [Boorla and Maranas, 2025], EITLEM-Kinetics [Shen et al., 2024], MPEK [Wang et al., 2024], CataPro [Wang et al., 2025] and DLKcat [Li et al., 2022] were evaluated on the same cleaned split definitions. DLKcat predicts *k*_cat_ only. The fixed-checkpoint curves report Spearman correlation with 95% confidence intervals obtained by bootstrapping test sequences. Fig. 3d reports the same fixed-checkpoint estimates at Seq-99, with 95% confidence intervals from the same sequence-bootstrap procedure. Performance was additionally summarized by *R*^2^, Pearson correlation, root-mean-square error and mean absolute error on the logarithmic targets. Complete fixed-checkpoint results are reported in Supplementary Tables S3 and S4.

### 4.6 Supervised within-landscape mutation-effect evaluation

The mutation model concatenated the 1,152-dimensional shared representation with a 1,152-dimensional raw sequence residual relative to the parent. The resulting 2,304-dimensional vector was standardized using training data and passed to a residual multilayer perceptron with hidden dimension 768, bottleneck dimension 384 and dropout 0.2. Training used 40 epochs, batch size 512, learning rate 2 × 10^−4^ and weight decay 8 × 10^−3^. Mean squared error was combined with a pairwise ranking term weighted by 0.05. Inputs excluded task-specific representations, mutation metadata and amino-acid one-hot encodings.

Evaluation used eight enzyme assays from ProteinGym [Notin et al., 2023]. We used eight leave-one-protein-out folds. In each fold, one protein was held out for testing and one for validation. The remaining six proteins were used for training. The eight test proteins cover all 41,165 single-substitution variants. Targets were standardized within assay using training-protein statistics. For each fold, eight pre-specified fusion candidates were evaluated under the same budget and selected by Spearman correlation on the validation protein, with nDCG@20 used to break ties. The selected fusion was then evaluated on the test protein. AlphaEnzyme scores are the average of three seeds.

Published baseline scores were aligned to the same test variants. For baselines with sign-ambiguous outputs, the training and validation proteins determined the score direction, which was then held fixed for testing. The fifteen external methods were the alignment-based models GEMME, DeepSequence and MSA Transformer [Laine et al., 2019, Riesselman et al., 2018, Rao et al., 2021]; the hybrid family-specific model TranceptEVE [Notin et al., 2022, Frazer et al., 2021]; the retrieval-conditioned language model PoET [Truong Jr. and Bepler, 2023]; the single-sequence language models ESM2 at 650M, 3B and 15B parameters, ESM-C and ESM3 [Lin et al., 2023, ESM Team, 2024, Hayes et al., 2025]; the structure-aware models SaProt, ProSST-2048, ProtSSN and ESM-IF1 [Su et al., 2024, Li et al., 2024, Tan et al., 2024, Hsu et al., 2022]; and the conservation-and-accessibility score RSALOR [Tsishyn et al., 2025]. Their model classes are shown in Fig. 3f, which displays a representative subset of the comparison. The complete 16-method comparison is reported in Supplementary Table S5. Metrics were calculated against the original ProteinGym DMS scores separately within each assay and then macro-averaged. They comprised Spearman and Pearson correlation, nDCG@20 and nDCG@50. For each assay and metric, methods displayed in Fig. 3f were ranked with ties assigned the minimum rank. Top-3 frequency reports the fraction of the resulting 32 assay–metric comparisons in which a method ranked first, second or third. Matched representation and WT-delta ablations are described in Supplementary Methods M2–M3 and Supplementary Fig. S9.

### 4.7 Retrospective TrpB prioritization

The TrpB data came from the four-site active-site fitness landscape of Johnston and colleagues [Johnston et al., 2024]. Fitness was the published growth-selection score, and parent-relative fitness was calculated against VFVS. Five disjoint 384-member libraries covered positions 183, 184, 227 and 228, with 128 double-, 128 triple- and 128 quadruple-substitution variants per library. Candidate identities were selected without fitness labels and fixed before AlphaEnzyme scoring. Supplementary Methods M5 details library construction and the separately trained reaction-aware TrpB readout.

AlphaEnzyme and the external models ranked the same candidates, with score directions fixed before label-based evaluation. The external score columns were EVmutation, PoET, SaProt, ESM3, ESM2-650M and ESM-C [Hopf et al., 2017, Truong Jr. and Bepler, 2023, Su et al., 2024, Hayes et al., 2025, Lin et al., 2023, ESM Team, 2024], alongside a random-selection reference. Model weights, hyperparameters and selection rules were fixed before the measurements were used for evaluation. This protocol evaluates transfer to the TrpB landscape without target-specific fitness supervision.

For a fixed budget *K*, the top *K* variants were selected within each library. Evaluation used the number of selected variants meeting the global Top-1% fitness criterion, their mean measured fitness and nDCG at *K*. Budget robustness was evaluated at *K* = 8, 12, 16, 20, 24, 28 and 32. Structural panels were prepared by mapping candidate side chains onto a fixed parent backbone around the pyridoxal 5′-phosphate cofactor. These models provided spatial context for the prioritized substitutions.

### 4.8 Candidate prioritization and experimental evaluation

Ten positions were predefined by combining structural context (catalytic pocket, PLP environment, flexible loops and second-shell residues), ESM-C single-substitution compatibility and structural-zone coverage [Höhne et al., 2010, Telzerow et al., 2021, Konia et al., 2021]. The selected positions were 58, 88, 126, 146, 183, 216, 235, 274, 290 and 309. Candidate construction used 16 allowed residue states per site and combined ESM-C-weighted proposals with uniform sampling at a ratio of 0.72:0.28 [Wittmann et al., 2021, Biswas et al., 2021]. Proposals covered two to ten selected sites across nine substitution-count layers and were deduplicated by mutation identity, giving a total collection of 10,000 sequences.

Representative selection grouped candidates by substitution count, one-hot encoded the selected residue states and applied MiniBatchKMeans clustering. The nearest observed sequence to each cluster centre was retained, with deduplication and filling to reach 1,000 representatives. The final panel was therefore a representative subset of the 10,000-member library. The ranking script applied the ProteinGym-trained mutation head described above and scored each candidate relative to the parent. The analysed experiment comprised 50 candidates from the AlphaEnzyme top-50 set, 20 candidates selected using ESM2 and 20 selected at random, with three biological replicate measurements per candidate. The rank-2 candidate carries substitutions I88D, Q216F, R235E, N290G and H309V relative to the 323-residue parent.

The target transformation was PLP-dependent transamination [Steffen-Munsberg et al., 2015, Schiroli and Peracchi, 2015] of 4-hydroxy-2-butanone with isopropylamine to (*R*)-3-aminobutan-1-ol (RABO) and acetone [Kelly et al., 2018, Patil et al., 2018] (Fig. 5b,c). Coding sequences were codon-optimized and synthesized, cloned into pET-28a under the T7 promoter and transformed into *Es-cherichia coli* C43(DE3) with kanamycin selection. Expression used 380 *μ*L Terrific Broth per well in 2.2 mL 96-deep-well plates at 30 ^◦^C, 800 rpm and 80% relative humidity, induced with 1 mM iso-propyl *β*-D-1-thiogalactopyranoside, and continued for 20 h. Pellets were frozen at −80 ^◦^C and resuspended in 100 mM triethanolamine·H_2_SO_4_ (pH 8.0) containing polymyxin B sulfate, lysozyme, DNase I and PLP. Lysates were clarified by centrifugation.

Biotransformations (200 *μ*L) combined 20 *μ*L clarified lysate with 180 *μ*L reaction buffer (1.2 M isopropylamine, 100 mg mL^−1^ 4-hydroxy-2-butanone and 0.081 g L^−1^ PLP in 100 mM triethanolamine·H_2_SO_4_, pH 8.0) and were incubated at 50 ^◦^C with shaking at 600 rpm for 20 h. A fixed lysate volume was used, so relative activity is expressed per unit lysate. Reactions were quenched with an equal volume of acetonitrile. Supernatants were derivatized with Marfey’s reagent (1-fluoro-2,4-dinitrophenyl-5-L-alaninamide, FDAA) [Marfey, 1984, Bhushan and Brückner, 2004] at 40 ^◦^C for 1 h and analysed on an Agilent 1290 Infinity II UHPLC with an EclipsePlus C18 RRHD column (2.1 × 50 mm, 1.8 *μ*m), isocratic water:acetonitrile (60:40, v/v) at 0.65 mL min^−1^, 40 ^◦^C and 340 nm over 2.5 min. Product identity and configuration were assigned against authentic (*R*)- and (*S*)-3-aminobutan-1-ol standards derivatized in parallel. The parent enzyme and the rank-2 variant both gave only the (*S*, *R*) adduct, consistent with exclusive formation of the (*R*) enantiomer (Supplementary Fig. S10). The full protocol is in Supplementary Methods M6.

The parent-normalized product signal was the RABO UPLC peak area divided by the mean parent peak area from the same plate. The parent mean was rounded to two decimal places before division. Means and sample standard deviations were calculated from the three biological replicate measurements per candidate. The rank-2 comparison used its own plate’s three parent controls. Recorded zeros indicate an undetected product signal. Absent candidate ranks were omitted rather than assigned zero. The displayed uncertainty summarizes biological-replicate variability. Measurements from the parallel campaign were not pooled with the current experiment.

### 4.9 Structural visualization of the experimentally prioritized variant

All structures were modelled with Protenix [ByteDance AML AI4Science Team et al., 2025], using one protein chain, PLP and 4-hydroxy-2-butanone. Alignment was reused with its query row replaced by each target sequence. Homologous sequences and alignment columns were retained. Predictions used three random seeds (101, 202 and 303), five diffusion samples per seed, ten recycling iterations and 200 diffusion steps, without structural templates or manually specified contacts.

Models were checked for sequence identity, backbone geometry and interchain clashes. The passing model with the highest Protenix ranking score was selected for each sequence. PyMOL rendered the coordinate-derived views with a pale-blue molecular surface, grey protein cartoon, blue design or substitution positions, orange PLP and teal substrate. These predicted complexes provide spatial context for the experimental variants. Selected coordinates, inputs and confidence records accompany the Fig. 5 source data.

## Data availability

Source-data tables accompanying the Supplementary Information provide the numerical results, evaluation records and analysis manifests used in the supplementary displays. Fig. 5 source data include the candidate measurements from the AlphaEnzyme top-50, ESM2 and random-selection sets, their model ranks and scores, individual UPLC measurements, parent controls and predicted complex coordinates.

## Code availability

The source code and reproducible inference entry points are available from https://github.com/ GENTEL-lab/AlphaEnzyme.

## Acknowledgements

This study was supported by the National Natural Science Foundation of China (62041209), the Natural Science Foundation of Shanghai (24ZR1440600), the Science and Technology Commission of Shanghai Municipality (24510714300), the Key project at CG level: the ability establishment of sustainable use for valuable Chinese medicine resources (2060302) and the Fundamental Research Funds for the Central Universities (N26JFB002). The transaminase engineering campaign was carried out in collaboration with Bota Biosciences (Hangzhou Enhe Biotechnology Co., Ltd.) under a joint research agreement with Shanghai Jiao Tong University, which designed and performed the experimental validation together with the Shanghai Jiao Tong University team.

## Author contributions

S.Z. and C.-T.C. conceived and supervised the project. Y.L., Z.S., R.S., Ji.W. and Y.H. implemented the models. Y.L., Z.S., Y.H., Ju.W. and Y.D. collected and curated the data. Y.L., Z.S., J.L. and S.Z. prepared the figures. J.L., R.S., Z.Z., B.H., X.L., H.L., J.C. and K.L. performed the experimental validation. Y.L., Z.S., J.L. and S.Z. wrote the manuscript. All authors discussed the results and approved the final manuscript.

