## Supplementary Information for "Harnessing catalytic knowledge to guide enzyme discovery and evolution with AlphaEnzyme"

The supplementary analyses follow the functional progression in the main Results: catalytic compatibility, quantitative catalytic properties and mutational response. Complete comparisons, controlled readouts and evaluation protocols support each functional level of the shared molecular representation. Source-data tables provide the records and exact numerical results underlying the supplementary figures.

#### Contents

Supplementary Methods S2
Figure S1. Shared representation and functional readouts S7
Figure S2. Evaluation coverage across functional scales S8
Figure S3. Query-resolved enzyme retrieval S9
Figure S4. Retrieval gains across joint-divergence strata S10
Figure S5. Specificity module ablation S11
Figure S6. Complete kinetic performance profiles S12
Figure S7. Measured and predicted kinetics on Seq-40 S13
Figure S8. Kinetic module ablation S14
Figure S9. Mutation representation and WT-delta ablation S15
Figure S10. Chiral analysis of the transaminase product S16

Table S1. Evaluation cohorts and mutation folds S17
Table S2. Complete retrieval metrics and paired comparisons S18
Tables S3–S4. Kinetic metrics S19
Table S5. Assay-resolved mutation correlations under protein-level holdout S21
Table S6. Mutation ablation under protein-level holdout S21
Supplementary References S22

### Supplementary Methods

#### M1. Representation and evaluation cohorts

AlphaEnzyme combines structural, reaction and protein/pocket context through three 384-dimensional projected components. Their concatenation defines the 1,152-dimensional representation, $\mathbf{h}_{\mathrm{AE}}$, used by the specificity, kinetic and mutation readouts (Supplementary Fig. S1). Structural inputs derive from the AlphaFold 3-style Protenix implementation[S1][S2]. Reaction descriptors use DRFP[S3], and pocket descriptors use P2Rank[S4]. The mutation readout additionally receives a 1,152-dimensional parent-relative sequence residual. Task-specific supervision is applied to the corresponding biochemical readout. The archived mutation feature bank used a zero-valued DRFP input and retained the fixed 1,152-dimensional output interface. The main mutation evaluation and three-seed input decomposition use this same bank.

The specificity collection comprised 478,356 training-partition and 25,414 validation-partition pairs assembled from RHEA[S5]; the active feature pipeline applied a 700-residue length limit. The time-separated evaluation followed the July 2023 annotation boundary. The EC-based retrieval comparison contains 104 reactions with complete four-level EC annotations, 5,589 candidate pairs and 231 positive annotations. Its EC-field scoring protocol is defined in the main Methods. The contextual HIT-EC reference averages three independently initialized models, trained on 136,059 EC-labelled sequences from the training partition; validation masked binary cross-entropy selected checkpoints within a five-epoch window. The broader reaction-to-enzyme comparison contains 279 common reaction queries with 515 positive annotations across all six methods. The matched-control experiment retains its archived 295-query evaluation cohort. Joint-divergence analyses in both cohorts contain 24 queries. These query universes are kept separate throughout the supplementary figures and tables. As in the main Methods, these evaluation cohorts are time-separated, and their pair labels remained outside gradient-based fitting.

The fixed-checkpoint kinetic analysis evaluates one Seq-99-trained checkpoint per model and target on nested Seq-40, Seq-60, Seq-80 and Seq-99 subsets. These thresholds describe maximum sequence identity to training. Protein sequences define the unit of resampling, while measurements define the unit of prediction. Fig. 3d shows these fixed-checkpoint estimates at Seq-99.

Mutation prediction uses eight ProteinGym assays[S6], comprising 41,165 single-substitution variants. We used eight leave-one-protein-out folds; in each fold, one protein was held out for testing and one for validation, and the remaining six proteins were used for training. The eight test proteins cover all 41,165 variants. Scores retain their original assay interpretation, including functional selection and activity-surrogate measurements. PTEN reports a growth-based phosphatase readout, and the RASH assay includes C-Raf binding/GEF-related function. The reported mutation metrics therefore refer to assay-specific fitness. Supplementary Table S1 gives the fold assignments, protein lengths and reference accessions.

#### M2. Module ablations

Specificity ablations used the fixed 295-query temporal control cohort and the same saved predictions. Full, w/o structure, w/o pocket and w/o reaction were compared. w/o structure removes structural single and pair inputs and the pocket branch; w/o pocket removes only the pocket branch; w/o reaction removes DRFP and ligand-associated structural entries. Bars are three-seed ensembles; open circles are individual seeds.

Kinetic ablations used the original fixed training and validation tables with three seeds, with model selection by validation RMSE. Full, w/o structure token, w/o pocket and w/o substrate were compared on the nested Seq-40, Seq-60, Seq-80 and Seq-99 subsets. w/o structure token removes the structure token but retains protein and pocket vectors; w/o pocket removes the dedicated pocket branch; w/o substrate removes the explicit substrate module. Retained branches may carry overlapping structural context.

Mutation ablations used eight leave-one-protein-out folds; six proteins were used for training, one for validation and one for testing. Full, w/o Alpha representation and w/o WT delta were compared. Each version used the same fusion candidates, selected on the validation protein by SCC (NDCG@20 tie-break), and the selected fusion was evaluated on the test protein. Scores are the average of three seeds. No new training was performed for the ablation.

#### M3. Kinetic and mutation readouts

The final kinetic readouts use neural TabM heads on the adapted $\mathbf{h}_{\mathrm{AE}}$ representation. Each head has four blocks, hidden width 768, ensemble size 32, dropout 0.1 and 2,986,016 parameters. Training uses mean squared error, batches of 512, learning rate $10^{-3}$, weight decay $10^{-3}$, gradient clipping at 5 and at most 150 epochs. Validation early stopping has patience 18; learning-rate reduction has patience 5 and a floor of $5 \times 10^{-5}$. The archived Seq-99 runs used fixed training and validation partitions with no record-ID overlap. Validation selected epochs 60 and 129. These are the fixed final-model predictions in Supplementary Figs. S6–S7; the module ablations in Fig. S8 use the separate protocol in M2.

The mutation head receives $[\mathbf{h}_{\mathrm{AE}}; \Delta\mathbf{h}_{\mathrm{seq}}]$ through a 2,304-dimensional interface. Input standardization and within-assay target standardization use the training proteins. A residual multilayer perceptron has hidden width 768, bottleneck width 384 and dropout 0.2. Training uses 40 epochs, batches of 512, learning rate $2 \times 10^{-4}$ and weight decay $8 \times 10^{-3}$. The loss combines mean squared error with a pairwise ranking term weighted by 0.05. For each fold, eight pre-specified fusion candidates were evaluated under the same budget and selected by Spearman correlation on the validation protein, with NDCG@20 used to break ties. AlphaEnzyme scores are the average of three seeds. The test protein was not used for fusion selection.

Published baseline mutation scores were aligned to the same test variants. When score direction is ambiguous, the training and validation proteins determined its sign before test evaluation. The comparison includes AlphaEnzyme and 15 archived external methods, including GEMME (main Fig. 3f; Supplementary Table S5). Rank-based metrics retain the tie ordering of the original evaluation. The matched representation and WT-delta ablations are reported in Supplementary Fig. S9 and Table S6.

#### M4. Metrics, uncertainty and source-data alignment

For reaction query $q$, binary relevance $r_{q,i}$ gives

$$\mathrm{nDCG@}K(q) = \frac{\sum_{i=1}^{K} r_{q,\pi(i)}/\log_2(i+1)}{\sum_{i=1}^{K} r_{q,\pi^*(i)}/\log_2(i+1)}, \qquad \text{(S1)}$$

where $\pi$ is the predicted ordering and $\pi^*$ is the ideal ordering. Top-$K$ success is the presence of at least one annotated positive among the first $K$ candidates. Enrichment factors compare the positive fraction in the selected top-ranked fraction with its prevalence in the full candidate set. Overall metrics are averaged over reaction queries. Sequence identity of 0.30 and reaction similarity of 0.60 define the four joint-divergence strata; equality belongs to the lower-similarity group.

Paired retrieval differences are formed on identical queries before bootstrap resampling. Query-level means and differences use 10,000 resamples within the relevant cohort or stratum. The 2.5th and 97.5th percentiles define the displayed pointwise 95% intervals. AlphaEnzyme stratum intervals in Table S2 retain the archived estimates used in main Fig. 2g; paired gains use the supplementary resampling protocol. For kinetic error summaries, 5,000 bootstrap replicates sample entire protein-sequence groups with replacement. All measurements of a sampled sequence remain together. Local reanalysis uses seed 20260903; the original training seeds and checkpoints are unchanged.

Kinetic errors are calculated in the original log-transformed target units, $\log_{10}(k_{\mathrm{cat}}/\mathrm{s}^{-1})$ or $\log_{10}(K_m/\mathrm{mM})$. Alongside $R^2$ and rank or linear correlation, we report root-mean-square and mean absolute error. We also report the fraction with absolute log error below 1, which corresponds to prediction within a factor of ten. Empirical error distributions retain every held-out measurement. Labels are aligned by record ID to the original strict-test tables to preserve evaluation precision; saved predictions are unchanged.

Mutation NDCG uses linear, non-negative relevance obtained by subtracting the minimum measured score within the evaluated assay from each fitness value, followed by the logarithmic discount above. Metrics are calculated per assay before equal weighting across the eight test proteins. Top-$K$ hit rate is the fraction of the selected variants whose measured fitness is at or above the 90th percentile of that assay; ties are included. The mutation ablation in Fig. S9 and Table S6 reports macro-averages over the eight test proteins. Source-data files retain sample identifiers, model names, condition labels, fold definitions and the values used in each display.

#### M5. TrpB candidate libraries and retrospective ranking

Five candidate libraries were drawn from one published TrpB landscape[S7]. For each construction seed, 10,000 variants were sampled using a mixture of ESM-C single-substitution priors (72%) and uniform sampling (28%). Pools contained 2,000 double, 4,000 triple and 4,000 quadruple substitutions relative to VFVS. Within each substitution-count group, MiniBatchKMeans clustering of one-hot site states selected 128 representative variants. Previously selected representatives were excluded, yielding five disjoint 384-member libraries and 1,920 unique candidates. Selection used seeds 20260803–20260807, and candidate identities were fixed before AlphaEnzyme scoring. Library manifests record the quotas and identity hashes.

The archived label-blind score matrix supplies AlphaEnzyme and six external score columns for the main comparison. Column identities, direction rules and available provenance are retained in the source-data inventory. Scores were frozen before fitness was joined for evaluation. The separately archived TrpB mutation readout uses the common representation interface with its reaction-aware input configuration. This linear readout was trained for eight epochs on the eight enzyme assays while the encoder remained fixed. Its training metadata, checkpoint identifier and feature settings are supplied with the source data. NDCG uses fitness shifted by the minimum within each candidate library. Global Top-1% membership is defined across the complete measured double-to-quadruple space, with variant-code ordering resolving tied boundary scores. In main Fig. 4b, whiskers show between-library s.d.; in Fig. 4c, ribbons show s.e.m. across the five libraries. All per-library metrics and all seven budgets are retained in the source data.

The global Top-1% criterion identifies 1,591 of the 159,052 measured double-to-quadruple variants in the published landscape. Random-selection references use 10,000 draws per library and budget. The reported 1.96-fold parent-relative value is calculated from the published growth-selection fitness of VIMG (0.800) and VFVS (0.408).

#### M6. Transaminase screening protocol

This section records the screening protocol for the transaminase campaign of main Fig. 5 at operational detail. The condensed description appears in Methods; the two are intended to be consistent, and this section is the reproducible reference.

*Constructs and host.* Coding sequences of the selected amine transaminases (ATAs) were codon-optimized and chemically synthesized. Each variant was cloned into the pET-28a vector under control of the T7 promoter and transformed into *Escherichia coli* C43(DE3) for cytoplasmic protein expression. Kanamycin was used as the selective antibiotic for both plasmid maintenance and expression cultures throughout the study.

*Cultivation and induction.* Pre-cultures were propagated in Luria–Bertani medium supplemented with kanamycin. For protein expression, 380 $\mu$L of kanamycin-containing Terrific Broth was inoculated per well in 2.2 mL 96-deep-well plates. Plates were incubated at 30 °C and 80% relative humidity with orbital shaking at 800 rpm for 2 h. Protein expression was induced by adding isopropyl $\beta$-D-1-thiogalactopyranoside to a final concentration of 1 mM, followed by further incubation at 30 °C, 800 rpm and 80% relative humidity overnight (20 h).

*Cell harvest and lysis.* Cultures were harvested by centrifugation (4,000 rpm, 4 °C, 20 min). Cell pellets were stored at −80 °C for a minimum of 60 min before lysis; this freeze–thaw treatment improves cell disruption efficiency. Lysis buffer was prepared immediately before use in 100 mM triethanolamine·$H_2SO_4$ buffer (pH 8.0; 14.919 g triethanolamine dissolved in 900 mL purified water, followed by addition of 100 mL 0.5 M $H_2SO_4$) containing 0.5 g $L^{-1}$ polymyxin B sulfate, 1 g $L^{-1}$ lysozyme, 0.05 g $L^{-1}$ pyridoxal 5′-phosphate (PLP) and 0.1% (v/v) DNase I. The PLP-supplemented buffer yielded a clear yellow solution. 400 $\mu$L lysis buffer was added to each well and pellets were resuspended by vortexing for 1 min. Plates were incubated at room temperature with shaking at 900 rpm for 90 min. After a short clarifying spin (4,000 rpm, 4 °C, 1 min), 400 $\mu$L of 0.05 g $L^{-1}$ PLP in triethanolamine buffer was dispensed into each well. Plates were shaken at 600 rpm for 10 min at room temperature, and clarified lysates were recovered by centrifugation at 4,000 rpm and 4 °C for 20 min.

*Biotransformation.* Reaction buffer was prepared freshly for each screening batch. For a total volume of 108 mL, 8.8 mg PLP was dissolved in 66 mL 100 mM triethanolamine·$H_2SO_4$ buffer (pH 8.0). This solution was combined with 30 mL of 4.8 M isopropylamine (IPM) stock solution (pH 9.71, verified immediately before use; prepared from 4.27 M isopropylamine hydrochloride and 0.53 M free isopropylamine) and 12 mL neat 4-hydroxy-2-butanone. The resulting mixture contained final concentrations of 0.081 g $L^{-1}$ PLP, 1.2 M IPM as amine donor and 100 mg $mL^{-1}$ ketone substrate. Reactions (200 $\mu$L total volume) were assembled in new 96-deep-well plates by transferring 20 $\mu$L clarified lysate supernatant with an automated liquid handler (i7), followed by addition of 180 $\mu$L reaction buffer. Plates were heat-sealed (165 °C, 2.5 s) and incubated at 50 °C with orbital shaking at 600 rpm for 20 h to convert 4-hydroxy-2-butanone to (*R*)-3-aminobutan-1-ol (RABO). A fixed lysate volume was dispensed rather than a normalized enzyme amount.

*Quenching and derivatization.* Reactions were quenched by addition of 200 $\mu$L acetonitrile. Mixtures were shaken at 600 rpm for 10 min at room temperature and centrifuged at 4,000 rpm for 10 min at room temperature. For Marfey's derivatization, 50 $\mu$L quenched supernatant was transferred to a deep-well plate pre-loaded with 50 $\mu$L 10% (w/v) $NaHCO_3$ solution. 250 $\mu$L Marfey's reagent (1-fluoro-2,4-dinitrophenyl-5-L-alaninamide, FDAA; 2 mg $mL^{-1}$ in acetonitrile, prepared immediately before use) was then added. Plates were sealed and incubated at 40 °C with shaking at 900 rpm for 1 h.

*UPLC analysis.* Derivatized samples were centrifuged (4,000 rpm, room temperature, 10 min) and 150 $\mu$L supernatant was transferred to a round-bottom plate. Analyses were performed on an Agilent 1290 Infinity II UHPLC system equipped with an Agilent EclipsePlus C18 RRHD column (2.1 × 50 mm, 1.8 $\mu$m). Mobile phase A was water and mobile phase B was acetonitrile. Separation used isocratic elution (A:B = 60:40, v/v) at a flow rate of 0.65 mL $min^{-1}$, a column temperature of 40 °C, a sample compartment temperature of 22 °C and an injection volume of 1 $\mu$L. The needle wash solvent was 50% acetonitrile and the seal wash solvent was 10% isopropanol. Marfey's adducts were detected by ultraviolet absorbance at 340 nm with a total run time of 2.5 min. The column itself is achiral; because FDAA is a chiral derivatizing agent, RABO and SABO form diastereomeric adducts that are separated under these conditions.

*Chiral assignment of the product.* The configuration of the enzymatic product was assigned by comparing derivatized reaction samples with a derivatized racemic ($R/S$)-3-aminobutan-1-ol reference standard and a blank reaction control analysed in the same sequence (Supplementary Fig. S10). The racemic standard resolved into two well-separated diastereomeric peaks, assigned as the ($S, S$) adduct formed from SABO (peak 1, earlier eluting) and the ($S, R$) adduct formed from RABO (peak 2, later eluting); neither peak was detected in the blank. Samples from both the parent enzyme and the rank-2 variant gave peak 2 only, with no detectable peak 1. The biotransformation therefore yields the ($R$)-configured amino alcohol, and the prioritized variant retains the ($R$)-selectivity of the parent.

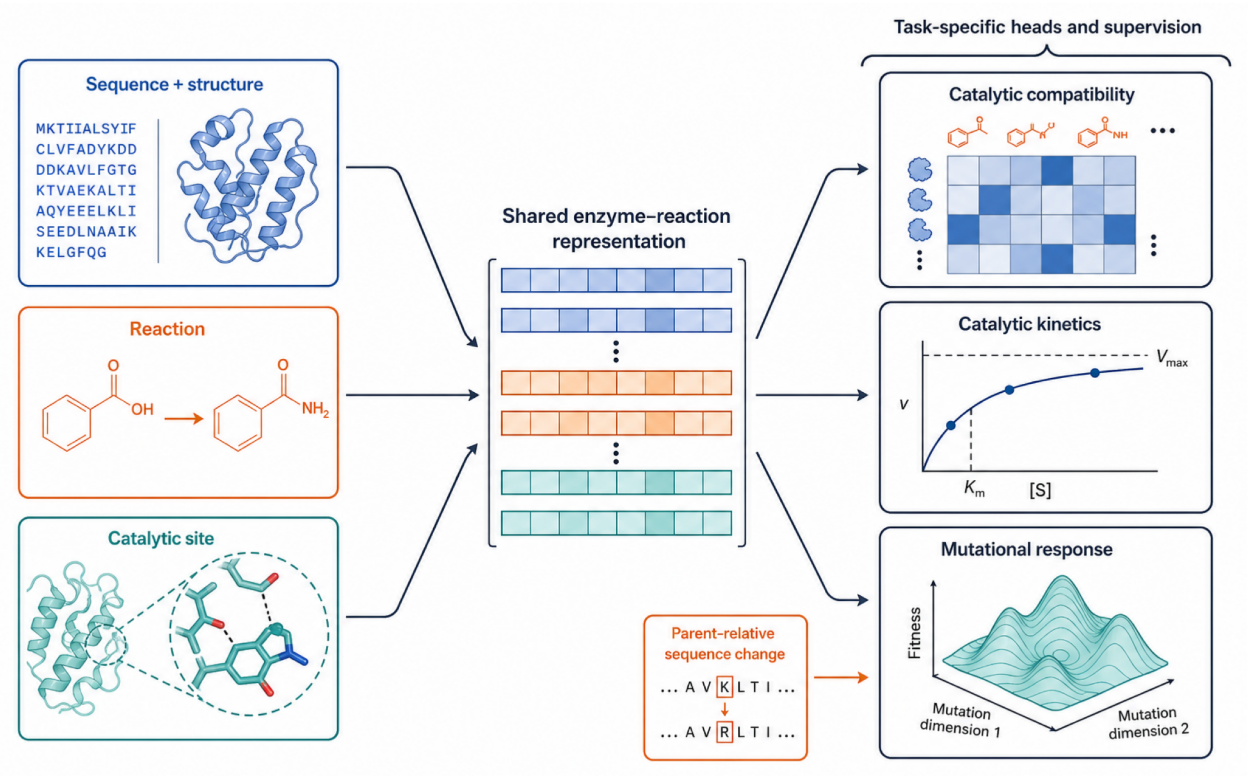


**Supplementary Figure S1. A shared enzyme–reaction representation supports task-specific functional readouts.** Sequence and structural context, reaction context and catalytic-site context are integrated into the shared AlphaEnzyme representation. Quantity-specific heads read from this common interface for catalytic compatibility, catalytic kinetics and mutational response. The mutation readout additionally receives the parent-relative sequence change. The three projected 384-dimensional blocks form a 1,152-dimensional shared representation. Addition of the sequence residual gives a 2,304-dimensional mutation input. The framework is shown in main Fig. 1; Supplementary Methods M1 and M3 define the exact inputs used in each evaluation.

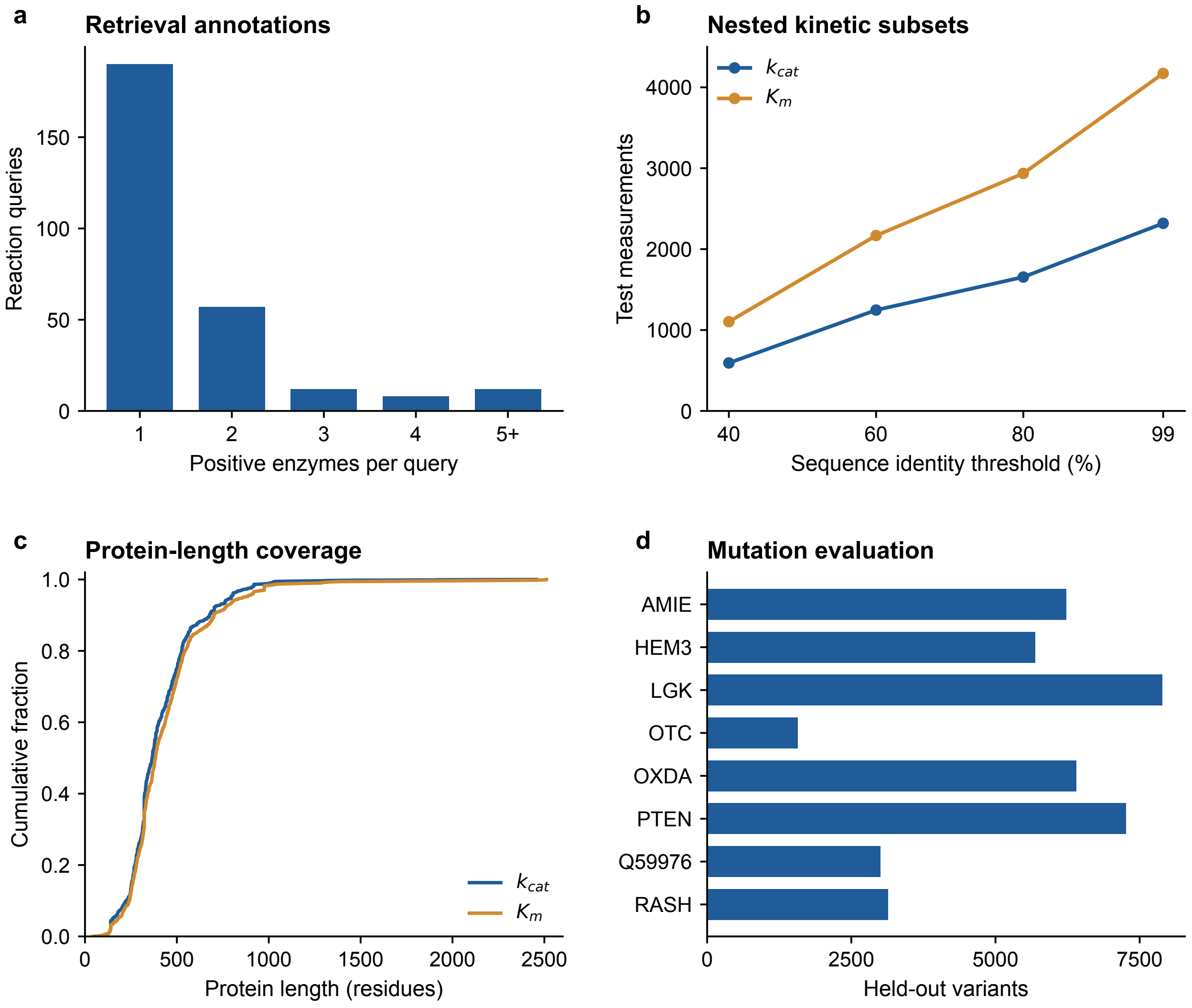


**Supplementary Figure S2. Evaluation coverage spans three linked functional scales. a**, Number of annotated positive enzymes per query in the common reaction-to-enzyme evaluation. The cohort contains 279 reaction queries and 515 positive relationships; queries with five or more positives are grouped in the rightmost bar. **b**, Numbers of $k_{\text{cat}}$ and $K_m$ measurements in the nested fixed-checkpoint evaluation, as the permitted maximum sequence identity to training increases from 40% to 99%. **c**, Protein-length distributions for the complete Seq-99 test sets. Each measurement contributes one observation. **d**, Held-out variant counts across the eight protein-level holdout folds; all 41,165 variants are covered across the eight folds (Table S1). Exact measurement counts and unique-sequence counts are retained in the source data; reference accessions and mutation folds are provided in Supplementary Table S1.

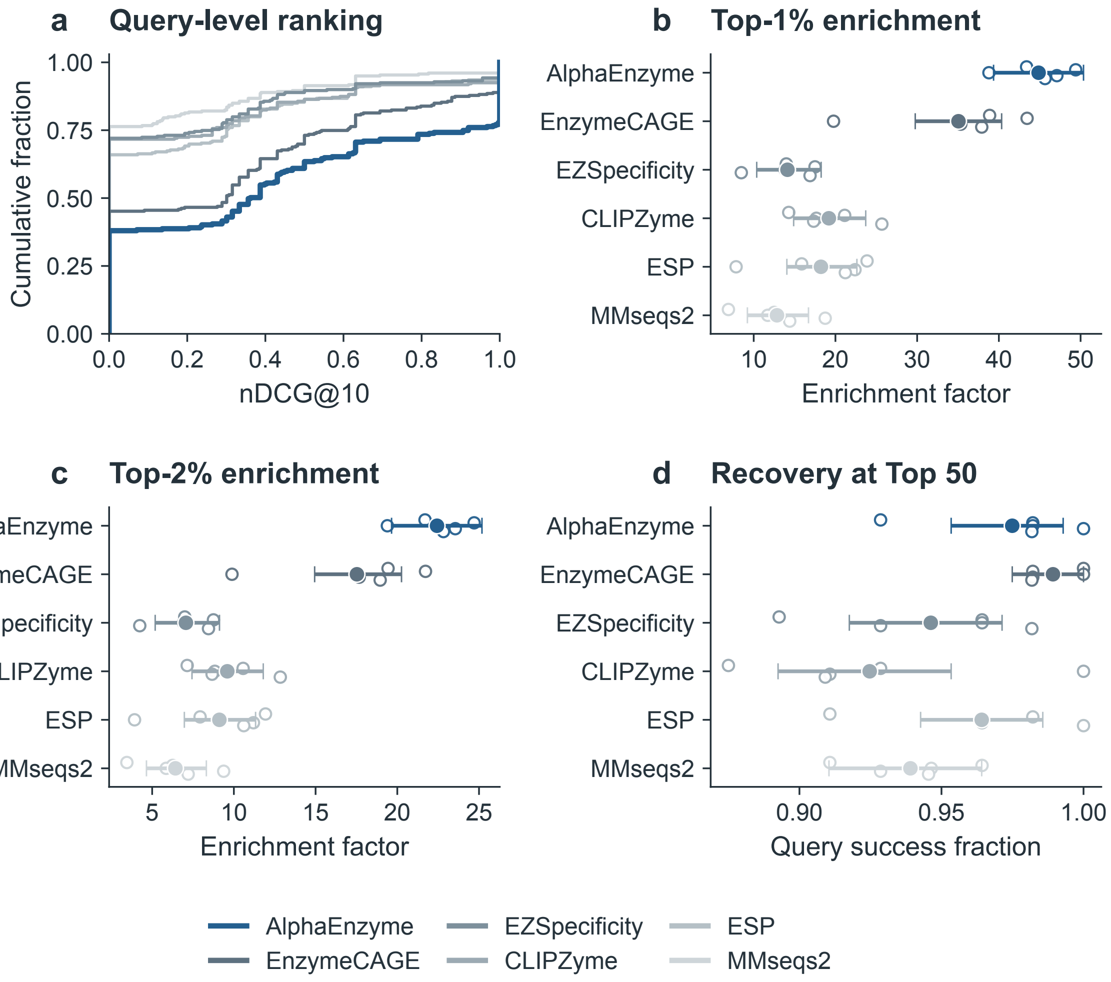


**Supplementary Figure S3. Query-level ranking and recovery across the enzyme-retrieval comparison.** **a**, Empirical cumulative distributions of query-level nDCG@10 for AlphaEnzyme and the five comparison methods. A shift towards larger nDCG values indicates stronger ranking of annotated catalysts. **b,c**, Mean enrichment factors among the top 1% and top 2% of ranked enzyme candidates. **d**, Fraction of queries with at least one known catalyst recovered among the top 50 candidates. In **b–d**, filled points show the full-query means, open points show five fixed query-fold estimates and whiskers show pointwise 95% query-bootstrap intervals from 10,000 resamples. All panels use the same 279 reaction queries and the same candidate sets across methods. The six methods are shown in a fixed order across panels; all known positives for each query contribute to the ranking metrics. Supplementary Table S2 reports Top-1, Top-3, Top-5, Top-10 and Top-50 success, enrichment factors, DCG@10 and nDCG@10.

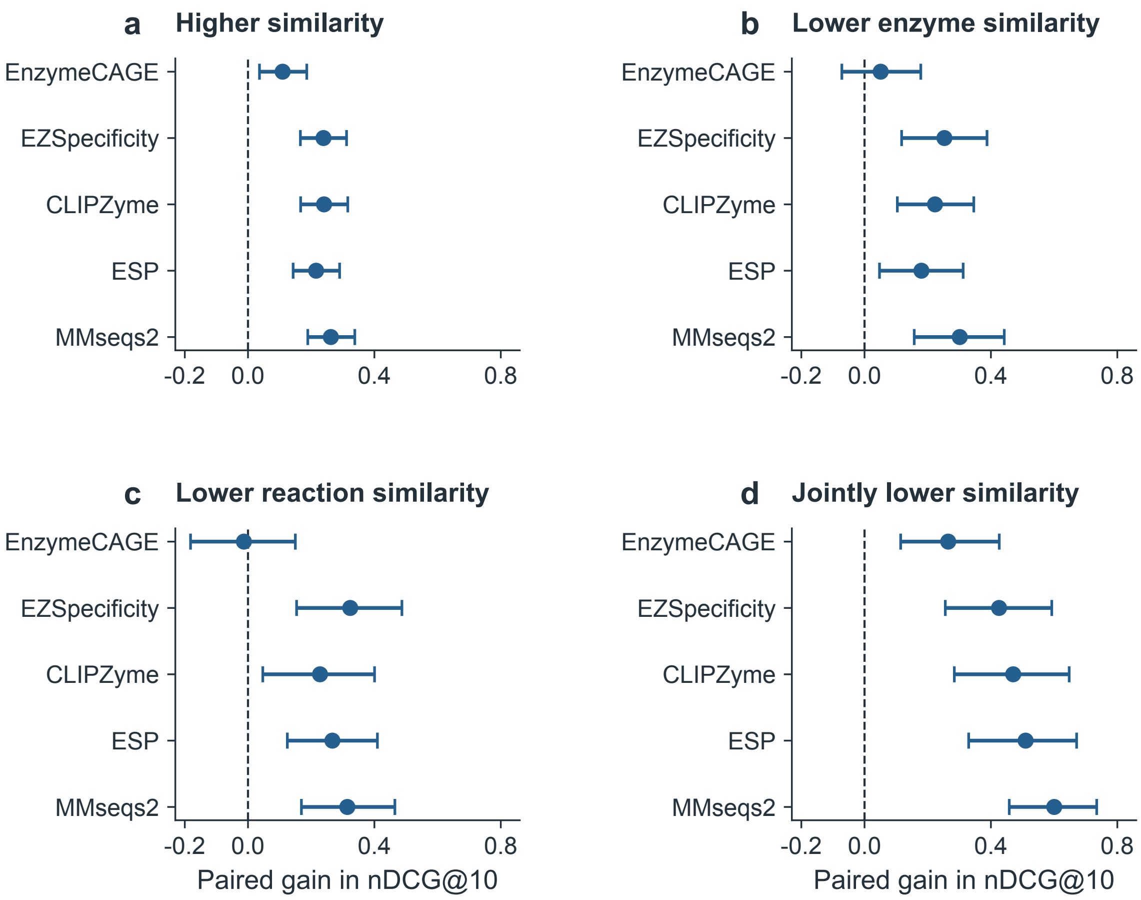


**Supplementary Figure S4. Paired retrieval gains persist under joint enzyme and reaction divergence.** Differences in query-level nDCG@10 between AlphaEnzyme and each comparison method, stratified by similarity to the specificity training data. **a**, Higher enzyme and reaction similarity ($n$ = 162). **b**, Lower enzyme similarity only ($n$ = 50). **c**, Lower reaction similarity only ($n$ = 43). **d**, Jointly lower enzyme and reaction similarity ($n$ = 24). Lower similarity is defined by maximum enzyme identity ≤ 0.30 or maximum reaction similarity ≤ 0.60. Points show the mean paired difference; whiskers show pointwise 95% percentile intervals from 10,000 within-stratum query-bootstrap resamples. Positive values favour AlphaEnzyme. In the jointly divergent stratum, its mean gain over EnzymeCAGE is 0.265 (95% interval, 0.114–0.426). All four strata and all five comparators are retained.

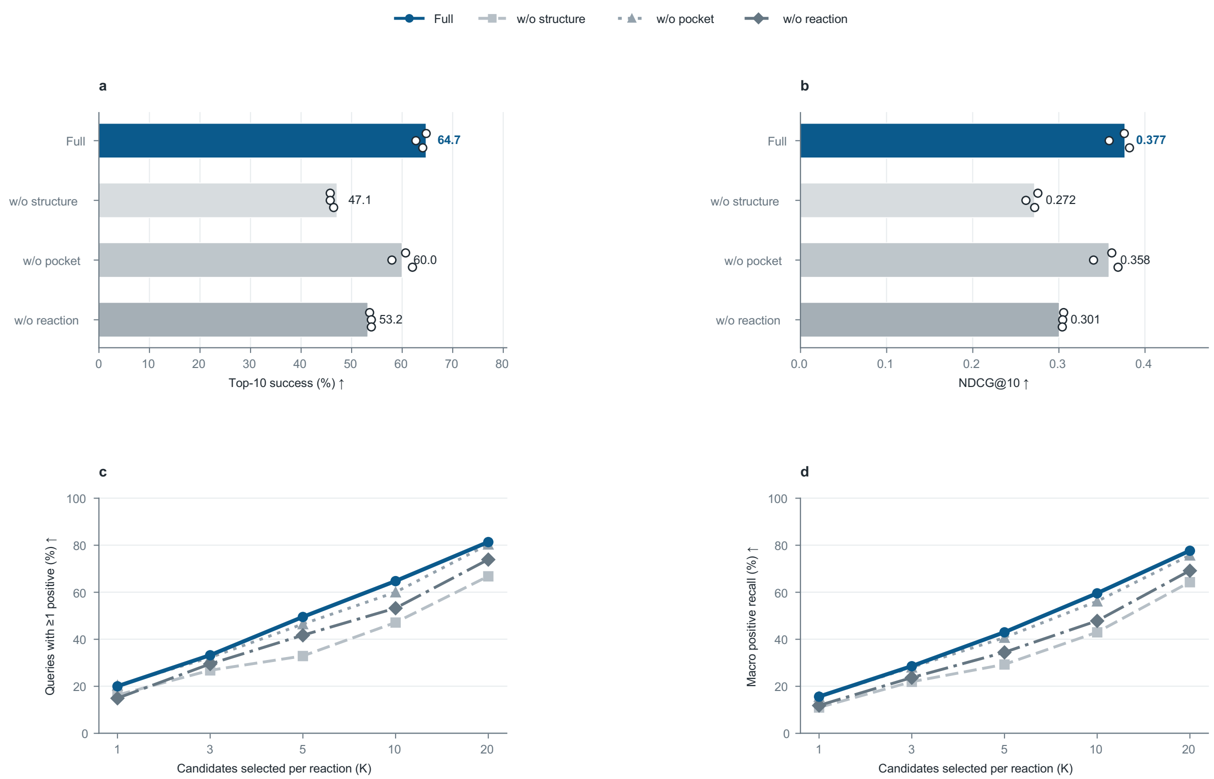


**Supplementary Figure S5. Specificity module ablation on the 295-query temporal control cohort. a**, Top-10 success and **b**, nDCG@10 for Full and three module ablations. **c**, Success and **d**, macro positive recall across selection budgets. Bars and curves are three-seed ensembles; open circles are individual seeds. Structure ablation removes the structural inputs and the pocket branch; pocket ablation removes only the pocket branch; reaction ablation removes DRFP and ligand-associated structural entries. This ablation uses the archived 295-query control cohort, so its Full condition is not numerically identical to the 279-query results in main Fig. 2 and Supplementary Table S2 (Methods M2).

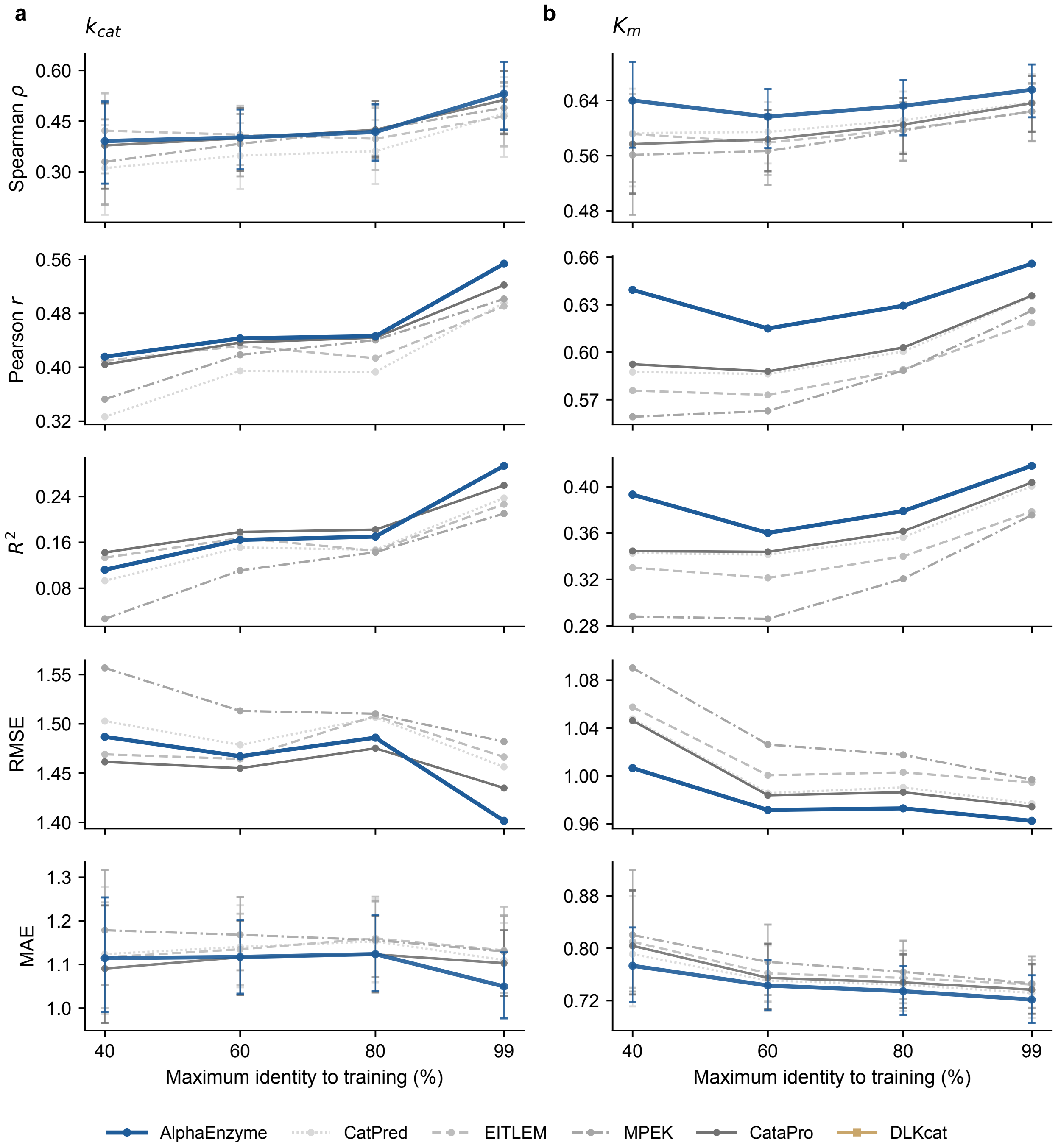


**Supplementary Figure S6. Quantitative catalytic information is retained across sequence-divergence levels.** Complete performance profiles for **a**, $k_{cat}$ and **b**, $K_m$ on the nested Seq-40, Seq-60, Seq-80 and Seq-99 test subsets. Rows show Spearman correlation, Pearson correlation, $R^2$, RMSE and MAE, respectively. AlphaEnzyme uses one fixed Seq-99-trained checkpoint across all four subsets; external methods use fixed pre-trained model weights. Error metrics use the log-transformed target units. Points mark the fixed-checkpoint estimates; whiskers shown for Spearman correlation and MAE are pointwise 95% bootstrap intervals obtained by resampling unique protein sequences. Exact measurement and unique-sequence counts are retained in the source data; all 44 model–target–threshold metric records are reported in Supplementary Tables S3 and S4. DLKcat is shown for $k_{cat}$ only.

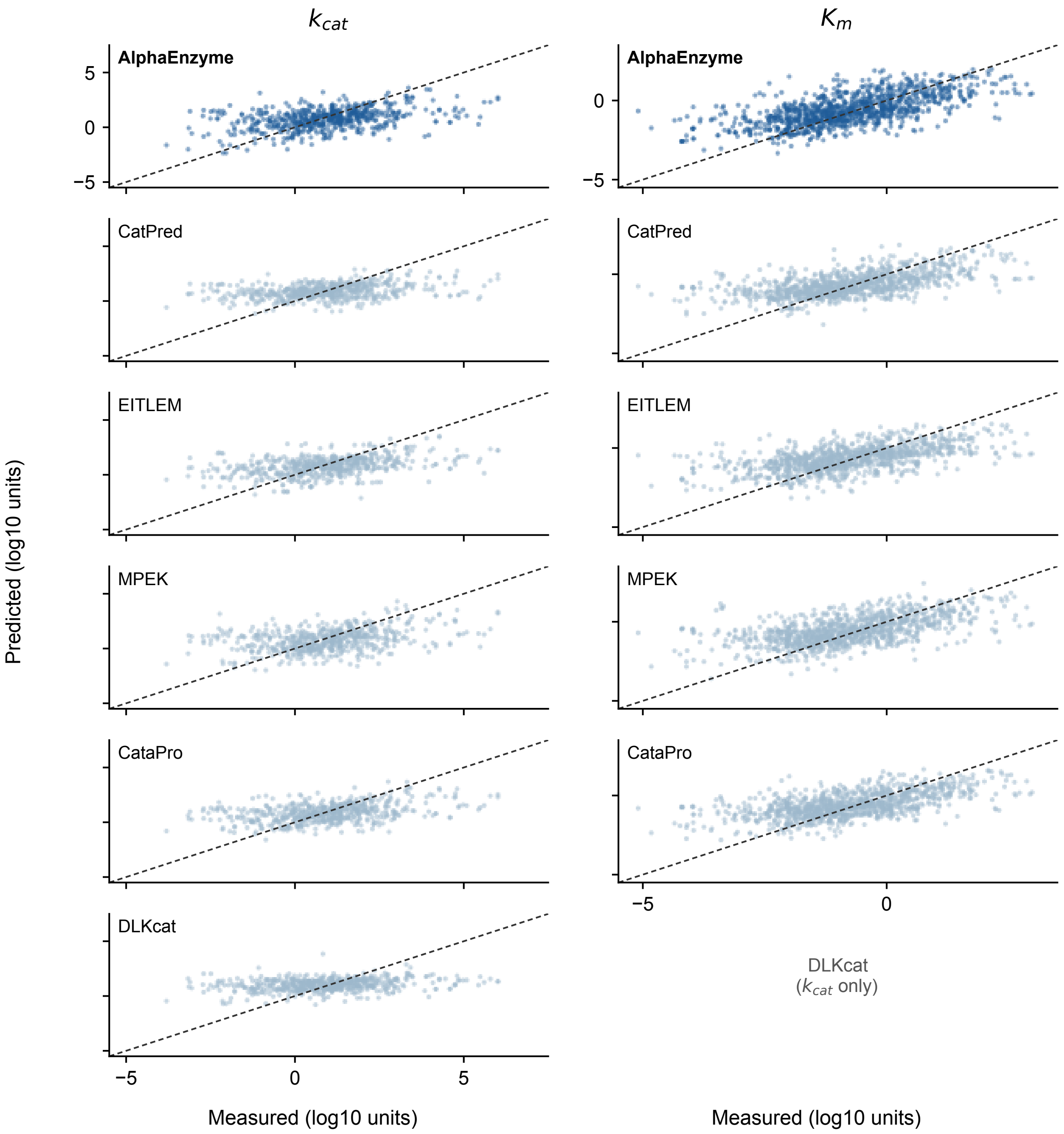


**Supplementary Figure S7. Measured and predicted catalytic parameters on the most sequence-divergent test subset.** Saved predictions for every Seq-40 measurement are plotted against the measured log-transformed target. Columns show $k_{\mathrm{cat}}$ and $K_m$; rows show AlphaEnzyme, CatPred, EITLEM, MPEK, CataPro and DLKcat. DLKcat predicts $k_{\mathrm{cat}}$ only; its $K_m$ panel is marked not applicable. Dashed lines indicate equal predicted and measured values. All methods use identical axis limits within each target and the same held-out records. The plots use the fixed Seq-99 checkpoints and retain every held-out measurement. Complete metrics accompany these predictions in Supplementary Tables S3 and S4.

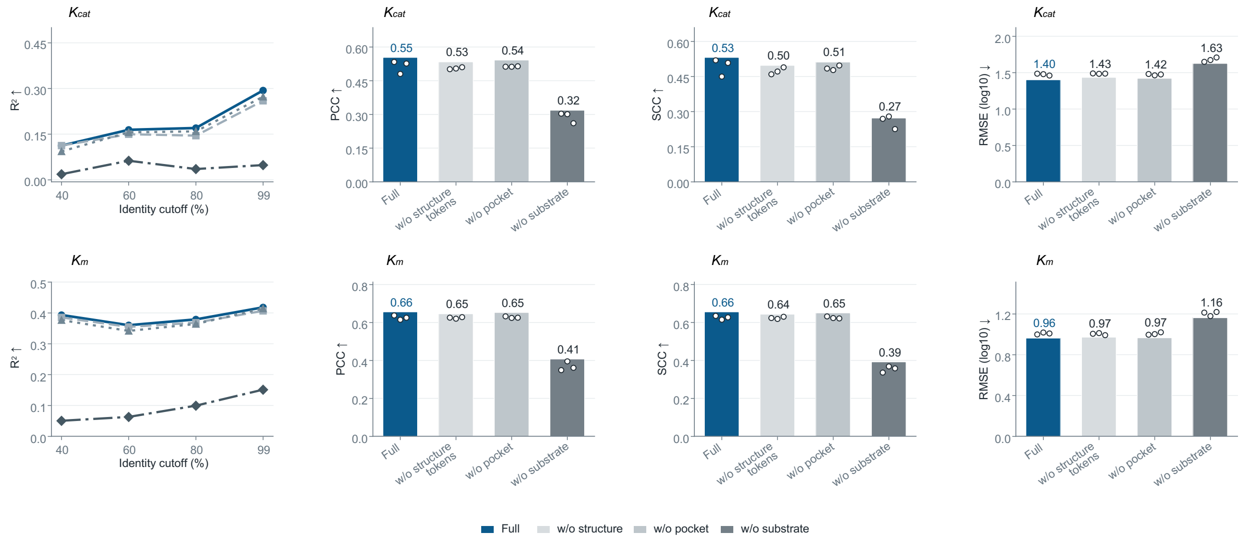


**Supplementary Figure S8. Kinetic prediction under structure-token, pocket and substrate ablations.** Rows show $k_{\text{cat}}$ and $K_m$. Columns show $R^2$ across Seq-40, Seq-60, Seq-80 and Seq-99, and PCC, SCC and RMSE at Seq-99, for Full and three ablations. Curves and bars are three-seed ensembles; open circles are individual seeds. Structure-token removal preserves pocket vectors; pocket removal preserves structure tokens; substrate removal removes the explicit substrate module.

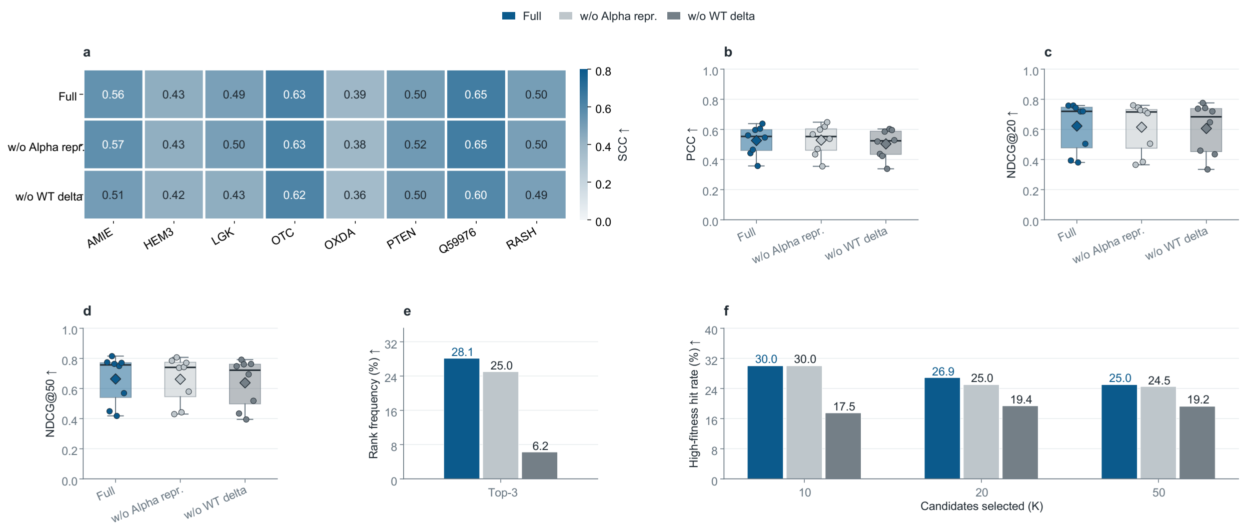


**Supplementary Figure S9. Mutation-effect prediction under representation and WT-delta ablations. a**, Per-protein SCC for Full and two ablations across eight protein-level holdouts. **b–d**, PCC, NDCG@20 and NDCG@50 distributions; points are proteins, diamonds are macro means. **e**, Top-3 rank frequency. **f**, High-fitness hit rate at $K = 10, 20, 50$ (fitness at or above each assay's 90th percentile). Full uses the validation-selected fusion; w/o Alpha removes the shared representation; w/o WT delta replaces the parent-relative difference with absolute mutant features. The Full condition is the validation-selected fusion and is therefore not numerically identical to the three-seed AlphaEnzyme ensemble reported in main Fig. 3f and Supplementary Table S5.

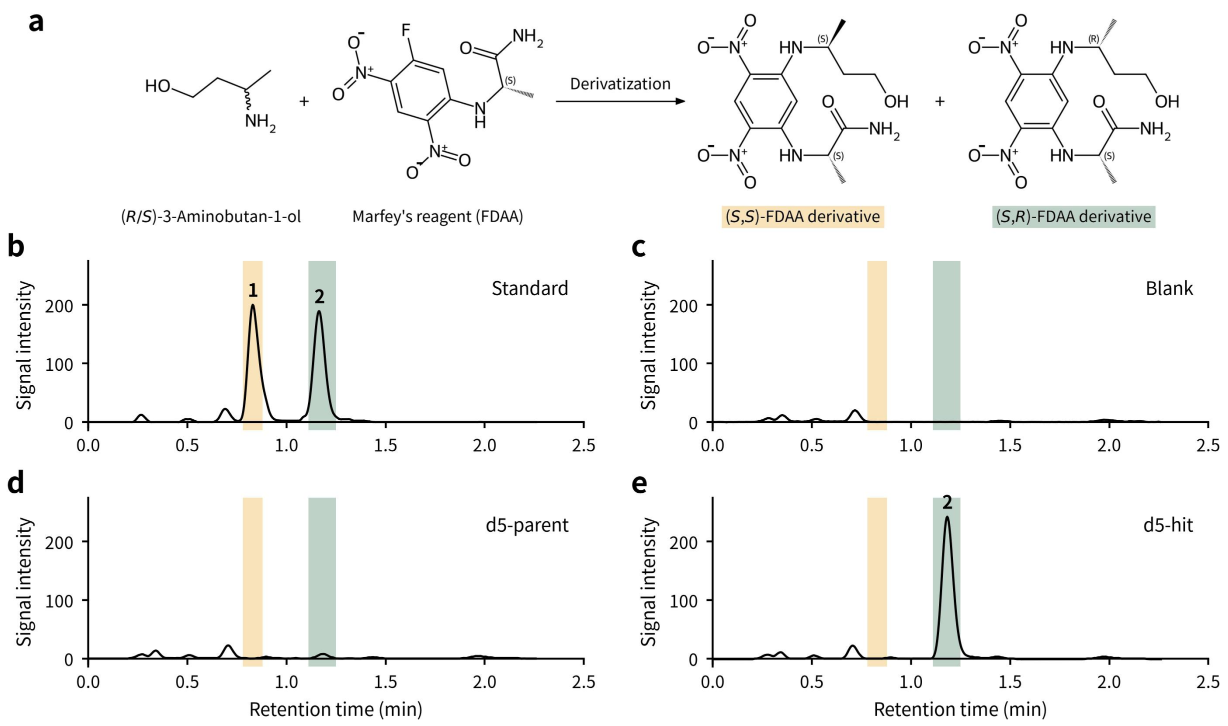


**Supplementary Figure S10. Chiral analysis of 3-aminobutan-1-ol by Marfey's FDAA derivatization coupled to UPLC.** Pre-column derivatization with Marfey's reagent (FDAA, 1-fluoro-2,4-dinitrophenyl-5-L-alaninamide) converts ($R$)- and ($S$)-3-aminobutan-1-ol into diastereomeric adducts that are separable on an achiral column; derivatization and chromatographic conditions are given in Supplementary Methods M6. **a**, FDAA derivatization of ($R/S$)-3-aminobutan-1-ol. The ($S,S$) adduct is formed from ($S$)-3-aminobutan-1-ol (SABO) and the ($S,R$) adduct from ($R$)-3-aminobutan-1-ol (RABO), the target product. **b–e**, UPLC chromatograms. Peak 1 corresponds to the ($S,S$)-FDAA derivative, and Peak 2 corresponds to the ($S,R$)-FDAA derivative. **b**, Racemic ($R/S$)-3-aminobutan-1-ol reference standard after derivatization, resolved into two well-separated diastereomeric peaks: peak 1, ($S,S$); peak 2, ($S,R$). **c**, Blank reaction control; no signal is detected in either peak window. **d**, Parent enzyme sample (parent). Only peak 2 is observed and peak 1 is absent, indicating that the product is the ($R$) enantiomer with no detectable ($S$)-configured product. **e**, Rank-2 variant sample (hit). Likewise only peak 2 is detected, with no observable peak 1, showing that the prioritized variant retains the stereoselective production of ($R$)-configured 3-aminobutan-1-ol. Signal intensity is plotted against retention time (min) in all chromatograms; shaded bands mark the peak 1 and peak 2 retention windows.

**Supplementary Table S1. Evaluation cohorts and exact assay partitions.** Counts describe the curated specificity partitions and the eight mutation folds. Kinetic subsets are evaluated in Fig. 3 and Tables S3–S4; the control-query cohort is maintained separately from the final-model comparison.

**a. Specificity learning and evaluation**

| Cohort | Count | Unit |
|---|---:|---|
| Curated training partition | 478,356 | Labelled pairs |
| Curated validation partition | 25,414 | Labelled pairs |
| EC-based retrieval comparison | 104 | Reaction queries |
| Final-model retrieval | 279 | Reaction queries |
| Positive retrieval annotations | 515 | Relationships |
| Matched specificity controls | 295 | Reaction queries |
| Jointly divergent subset | 24 | Reaction queries |

**b. Mutation evaluation folds**

| Fold | Test protein | Validation protein | Variants |
|---|---|---|---:|
| 0 | AMIE (P11436) | HEM3 | 6,227 |
| 1 | HEM3 (P08397) | LGK | 5,689 |
| 2 | LGK (B3VI55) | OTC | 7,890 |
| 3 | OTC (P00480) | OXDA | 1,570 |
| 4 | OXDA (P80324) | PTEN | 6,396 |
| 5 | PTEN (O00633) | Q59976 | 7,260 |
| 6 | Q59976 | RASH | 2,999 |
| 7 | RASH (P01112) | AMIE | 3,134 |
| Total | – | – | 41,165 |

All 41,165 mutation records carry one substitution. Mutation evaluation uses eight leave-one-protein-out folds; each fold holds out one protein for testing and one for validation, with the remaining six proteins used for training. Metrics are macro-averaged over the eight test proteins. Protein lengths and reference accessions are retained in the source data.

**Supplementary Table S2. Complete reaction-to-enzyme retrieval results.** The EC-based comparison uses 104 reactions and 5,589 candidate pairs. The broader comparison uses 279 reactions; its four similarity strata are evaluated without retraining.

**a. EC-based retrieval on 104 reaction queries**

| Method | Top-1 | Top-3 | Top-5 | Top-10 | DCG@10 | nDCG@10 |
|---|---|---|---|---|---|---|
| AlphaEnzyme | 19.2 | 28.8 | 38.5 | 53.8 | 0.438 | 0.307 |
| CLEAN | 14.4 | 15.4 | 18.3 | 22.1 | 0.261 | 0.157 |
| GraphEC | 6.7 | 8.7 | 16.3 | 20.2 | 0.198 | 0.122 |
| TopEC | 6.7 | 15.4 | 23.1 | 32.7 | 0.208 | 0.154 |
| HIT-EC | 12.5 | 18.3 | 24.7 | 34.9 | 0.276 | 0.179 |

The EC cohort contains 231 positive pairs. The contextual HIT-EC reference averages three retrained seeds.

**b. Full-cohort ranking and recovery on 279 reaction queries**

| Method | Top-1 | Top-3 | Top-5 | Top-10 | Top-50 | EF1 | EF2 | DCG | nDCG |
|---|---|---|---|---|---|---|---|---|---|
| AlphaEnzyme | 30.1 | 40.9 | 49.5 | 62.0 | 97.5 | 44.86 | 22.43 | 0.563 | 0.421 |
| EnzymeCAGE | 17.2 | 31.9 | 39.4 | 54.8 | 98.9 | 35.06 | 17.53 | 0.471 | 0.327 |
| EZSpecificity | 8.2 | 12.9 | 17.2 | 28.0 | 94.6 | 14.13 | 7.07 | 0.246 | 0.150 |
| CLIPZyme | 9.7 | 14.7 | 23.3 | 28.3 | 92.5 | 19.20 | 9.60 | 0.239 | 0.165 |
| ESP | 10.4 | 17.9 | 22.9 | 34.1 | 96.4 | 18.23 | 9.11 | 0.250 | 0.178 |
| MMseqs2 | 5.4 | 13.6 | 16.1 | 23.7 | 93.9 | 12.85 | 6.42 | 0.149 | 0.114 |

Top-$K$: success percentage; EF1/EF2: enrichment at the top 1%/2%. DCG and nDCG use rank 10.

**c. Mean nDCG@10 [95% interval] within each stratum**

| Method | Higher similarity | Lower enzyme | Lower reaction | Jointly lower |
|---|---|---|---|---|
| AlphaEnzyme | 0.379 [0.320, 0.438] | 0.416 [0.306, 0.528] | 0.452 [0.324, 0.589] | 0.657 [0.527, 0.785] |
| EnzymeCAGE | 0.269 [0.218, 0.321] | 0.365 [0.263, 0.472] | 0.465 [0.360, 0.572] | 0.392 [0.239, 0.544] |
| EZSpecificity | 0.140 [0.100, 0.183] | 0.163 [0.083, 0.255] | 0.129 [0.056, 0.218] | 0.231 [0.102, 0.374] |
| CLIPZyme | 0.138 [0.097, 0.183] | 0.193 [0.106, 0.286] | 0.225 [0.126, 0.333] | 0.186 [0.068, 0.325] |
| ESP | 0.163 [0.120, 0.208] | 0.236 [0.143, 0.339] | 0.185 [0.100, 0.282] | 0.147 [0.064, 0.253] |
| MMseqs2 | 0.117 [0.081, 0.155] | 0.115 [0.045, 0.198] | 0.138 [0.059, 0.227] | 0.056 [0.007, 0.115] |

**d. Paired AlphaEnzyme gain [95% interval]**

| Comparator | Higher similarity | Lower enzyme | Lower reaction | Jointly lower |
|---|---|---|---|---|
| EnzymeCAGE | 0.110 [0.036, 0.186] | 0.051 [-0.072, 0.178] | -0.013 [-0.182, 0.150] | 0.265 [0.114, 0.426] |
| EZSpecificity | 0.239 [0.166, 0.312] | 0.253 [0.117, 0.388] | 0.324 [0.154, 0.487] | 0.426 [0.256, 0.593] |
| CLIPZyme | 0.241 [0.167, 0.316] | 0.223 [0.104, 0.346] | 0.228 [0.047, 0.401] | 0.470 [0.284, 0.648] |
| ESP | 0.216 [0.143, 0.290] | 0.180 [0.047, 0.312] | 0.267 [0.125, 0.410] | 0.510 [0.330, 0.671] |
| MMseqs2 | 0.262 [0.189, 0.338] | 0.301 [0.157, 0.442] | 0.314 [0.169, 0.465] | 0.600 [0.458, 0.735] |

Stratum sizes are 162, 50, 43 and 24. Lower enzyme identity is ≤ 0.30; lower reaction-fingerprint similarity is ≤ 0.60. Intervals use 10,000 query-bootstrap resamples; paired gains use matched queries. Positive gains favour AlphaEnzyme.

**Supplementary Table S3. Complete fixed-checkpoint $k_{cat}$ evaluation.** All methods use the same fixed-checkpoint test records and three-seed prediction ensembles; DLKcat predicts $k_{cat}$ only.

| Seq. | Method | $R^2$ | $\rho$ | $r$ | RMSE | MAE | $P_{10}$ (%) |
|---|---|---|---|---|---|---|---|
| 40 | CatPred | 0.093 | 0.312 | 0.327 | 1.503 | 1.124 | 56.5 |
| 40 | EITLEM | 0.133 | 0.422 | 0.410 | 1.469 | 1.117 | 55.3 |
| 40 | MPEK | 0.027 | 0.330 | 0.353 | 1.557 | 1.179 | 55.6 |
| 40 | CataPro | 0.142 | 0.378 | 0.404 | 1.461 | 1.091 | 57.7 |
| 40 | DLKcat | 0.058 | 0.248 | 0.253 | 1.532 | 1.165 | 55.0 |
| 40 | AlphaEnzyme | 0.112 | 0.391 | 0.416 | 1.487 | 1.114 | 57.5 |
| 60 | CatPred | 0.151 | 0.348 | 0.395 | 1.479 | 1.140 | 53.3 |
| 60 | EITLEM | 0.167 | 0.410 | 0.432 | 1.464 | 1.135 | 52.9 |
| 60 | MPEK | 0.111 | 0.383 | 0.418 | 1.513 | 1.168 | 52.3 |
| 60 | CataPro | 0.178 | 0.399 | 0.437 | 1.455 | 1.116 | 54.5 |
| 60 | DLKcat | 0.096 | 0.297 | 0.311 | 1.526 | 1.179 | 52.1 |
| 60 | AlphaEnzyme | 0.164 | 0.402 | 0.443 | 1.467 | 1.117 | 55.8 |
| 80 | CatPred | 0.147 | 0.361 | 0.393 | 1.507 | 1.153 | 53.5 |
| 80 | EITLEM | 0.145 | 0.398 | 0.414 | 1.508 | 1.160 | 52.9 |
| 80 | MPEK | 0.143 | 0.425 | 0.441 | 1.510 | 1.156 | 53.2 |
| 80 | CataPro | 0.182 | 0.425 | 0.444 | 1.475 | 1.123 | 54.7 |
| 80 | DLKcat | 0.088 | 0.284 | 0.298 | 1.558 | 1.202 | 51.9 |
| 80 | AlphaEnzyme | 0.170 | 0.418 | 0.446 | 1.486 | 1.124 | 55.8 |
| 99 | CatPred | 0.237 | 0.472 | 0.496 | 1.456 | 1.110 | 55.6 |
| 99 | EITLEM | 0.227 | 0.465 | 0.491 | 1.466 | 1.132 | 53.8 |
| 99 | MPEK | 0.210 | 0.489 | 0.501 | 1.482 | 1.130 | 55.3 |
| 99 | CataPro | 0.260 | 0.513 | 0.522 | 1.435 | 1.103 | 55.0 |
| 99 | DLKcat | 0.095 | 0.304 | 0.320 | 1.586 | 1.241 | 49.6 |
| 99 | AlphaEnzyme | 0.294 | 0.531 | 0.554 | 1.402 | 1.050 | 59.0 |

$\rho$ and $r$ denote Spearman and Pearson correlation. $P_{10}$ is the fraction with absolute log error below 1. DLKcat predicts $k_{cat}$ only and is evaluated on the same test records.

**Supplementary Table S4. Complete fixed-checkpoint $K_m$ evaluation.** All methods use the same fixed-checkpoint test records and three-seed prediction ensembles.

| Seq. | Method | $R^2$ | $\rho$ | $r$ | RMSE | MAE | $P_{10}$ (%) |
|---|---|---|---|---|---|---|---|
| 40 | CatPred | 0.343 | 0.593 | 0.588 | 1.048 | 0.791 | 70.4 |
| 40 | EITLEM | 0.330 | 0.592 | 0.576 | 1.057 | 0.810 | 68.3 |
| 40 | MPEK | 0.288 | 0.561 | 0.559 | 1.090 | 0.820 | 69.0 |
| 40 | CataPro | 0.344 | 0.577 | 0.592 | 1.046 | 0.804 | 68.9 |
| 40 | AlphaEnzyme | 0.393 | 0.640 | 0.639 | 1.007 | 0.773 | 70.1 |
| 60 | CatPred | 0.341 | 0.594 | 0.586 | 0.986 | 0.751 | 72.9 |
| 60 | EITLEM | 0.321 | 0.579 | 0.573 | 1.000 | 0.762 | 71.9 |
| 60 | MPEK | 0.286 | 0.567 | 0.563 | 1.026 | 0.779 | 71.5 |
| 60 | CataPro | 0.344 | 0.584 | 0.588 | 0.984 | 0.755 | 72.9 |
| 60 | AlphaEnzyme | 0.360 | 0.616 | 0.615 | 0.971 | 0.743 | 72.3 |
| 80 | CatPred | 0.356 | 0.611 | 0.600 | 0.990 | 0.744 | 73.3 |
| 80 | EITLEM | 0.340 | 0.598 | 0.589 | 1.003 | 0.755 | 72.3 |
| 80 | MPEK | 0.321 | 0.597 | 0.588 | 1.018 | 0.764 | 72.5 |
| 80 | CataPro | 0.362 | 0.605 | 0.603 | 0.986 | 0.748 | 73.0 |
| 80 | AlphaEnzyme | 0.379 | 0.632 | 0.629 | 0.973 | 0.734 | 73.1 |
| 99 | CatPred | 0.400 | 0.637 | 0.635 | 0.977 | 0.732 | 74.2 |
| 99 | EITLEM | 0.378 | 0.624 | 0.619 | 0.995 | 0.745 | 73.4 |
| 99 | MPEK | 0.376 | 0.624 | 0.626 | 0.997 | 0.746 | 73.8 |
| 99 | CataPro | 0.404 | 0.636 | 0.636 | 0.974 | 0.737 | 74.4 |
| 99 | AlphaEnzyme | 0.418 | 0.656 | 0.656 | 0.962 | 0.722 | 74.2 |

$\rho$ and $r$ denote Spearman and Pearson correlation. $P_{10}$ is the fraction with absolute log error below 1.

**Supplementary Table S5. Assay-resolved Spearman correlation under protein-level holdout.** Values use the same eight protein-level test sets and 15 external methods. AlphaEnzyme is the three-seed ensemble; external entries are published scores aligned to the same test proteins. Mean is the equal-weight average of the eight displayed values.

| Method | AMIE | HEM3 | LGK | OTC | OXDA | PTEN | Q59976 | RASH | Mean |
|---|---|---|---|---|---|---|---|---|---|
| AlphaEnzyme | 0.56 | 0.44 | 0.50 | 0.64 | 0.41 | 0.50 | 0.65 | 0.50 | 0.525 |
| GEMME | 0.56 | 0.43 | 0.54 | 0.60 | 0.38 | 0.53 | 0.68 | 0.43 | 0.519 |
| TranceptEVE | 0.48 | 0.44 | 0.54 | 0.60 | 0.36 | 0.53 | 0.67 | 0.49 | 0.514 |
| RSALOR | 0.58 | 0.43 | 0.52 | 0.54 | 0.42 | 0.52 | 0.64 | 0.45 | 0.513 |
| PoET | 0.65 | 0.41 | 0.46 | 0.53 | 0.36 | 0.56 | 0.59 | 0.50 | 0.507 |
| SaProt | 0.61 | 0.43 | 0.41 | 0.62 | 0.40 | 0.54 | 0.63 | 0.37 | 0.501 |
| ProSST-2048 | 0.49 | 0.38 | 0.46 | 0.62 | 0.36 | 0.51 | 0.57 | 0.62 | 0.501 |
| ESM2-650M | 0.56 | 0.39 | 0.52 | 0.53 | 0.37 | 0.52 | 0.57 | 0.50 | 0.495 |
| ESM3 | 0.47 | 0.43 | 0.43 | 0.67 | 0.34 | 0.54 | 0.58 | 0.46 | 0.490 |
| ESM2-3B | 0.67 | 0.42 | 0.58 | 0.51 | 0.39 | 0.31 | 0.56 | 0.44 | 0.485 |
| ProtSSN | 0.54 | 0.37 | 0.52 | 0.57 | 0.37 | 0.51 | 0.54 | 0.43 | 0.481 |
| ESM-C | 0.50 | 0.40 | 0.41 | 0.58 | 0.38 | 0.54 | 0.66 | 0.37 | 0.480 |
| DeepSequence | 0.50 | 0.40 | 0.48 | 0.48 | 0.36 | 0.51 | 0.63 | 0.44 | 0.475 |
| MSA Transformer | 0.40 | 0.43 | 0.47 | 0.59 | 0.28 | 0.50 | 0.67 | 0.43 | 0.471 |
| ESM2-15B | 0.61 | 0.43 | 0.58 | 0.53 | 0.40 | 0.29 | 0.57 | 0.31 | 0.465 |
| ESM-IF1 | 0.41 | 0.30 | 0.49 | 0.67 | 0.36 | 0.48 | 0.53 | 0.46 | 0.463 |

Values are per-protein Spearman correlations under eight leave-one-protein-out folds. AlphaEnzyme is the three-seed ensemble; external methods are published scores aligned to the same test proteins. Means are equal-weight averages of the displayed two-decimal values. The comparison includes 15 external methods. Unrounded values and external source scores are retained in the accompanying source data.

**Supplementary Table S6. Mutation ablation under eight protein-level holdouts.** Macro-averages over the eight test proteins. Full uses the validation-selected fusion; w/o Alpha removes the shared representation; w/o WT delta replaces the parent-relative difference with absolute mutant features. Top-3 frequency is across 32 assay–metric comparisons.

| Condition | NDCG@20 | NDCG@50 | Top-10 | Top-20 | Top-50 | Top-3 |
|---|---|---|---|---|---|---|
| Full | 0.622 | 0.664 | 0.300 | 0.269 | 0.250 | 0.281 |
| w/o Alpha repr. | 0.615 | 0.662 | 0.300 | 0.250 | 0.245 | 0.250 |
| w/o WT delta | 0.606 | 0.638 | 0.175 | 0.194 | 0.193 | 0.062 |

Top-*K* columns are the fractions of selected variants whose measured fitness is at or above the assay-specific 90th percentile. Top-3 is the fraction of 32 assay–metric comparisons in which a condition ranked among the three highest-performing models of the mutation comparison displayed in main Fig. 3f.
